# Type I IFN signaling mediates NET release to promote *Mycobacterium tuberculosis* replication and granuloma caseation

**DOI:** 10.1101/2022.11.29.518376

**Authors:** Chanchal Sur Chowdhury, Rachel L. Kinsella, E. Michael Nehls, Sumanta K. Naik, Daniel S. Lane, Priyanka Talukdar, Sthefany M. Chavez, Asya Smirnov, Wandy Beatty, Darren Kreamalmeyer, Joshua T. Mattila, Christina L. Stallings

## Abstract

Neutrophils are the most abundant cell type in airways of tuberculosis patients. Recent investigations reported induction of neutrophil extracellular traps (NETs) during *Mycobacterium tuberculosis* (*Mtb*) infection, however, the molecular regulation and impact of NETosis on *Mtb* pathogenesis is unknown. We find that in response to *Mtb* infection in neutrophils, PAD4 citrullinates histones to decondense chromatin that gets packaged into vesicles for release as NETs in a manner that can maintain neutrophil viability and promote *Mtb* replication. Type I interferon, which has been associated with NETosis in numerous contexts but without a known mechanism, promotes formation of chromatin-containing vesicles and NET release. Analysis of nonhuman primate granulomas supports a model where neutrophils are exposed to type I interferon from macrophages as they migrate into the granuloma, where they release NETs that contribute to necrosis and caseation. Our data reveals NETosis as a promising target to inhibit *Mtb* replication and granuloma caseation.

## INTRODUCTION

Tuberculosis (TB) remains a leading cause of death due to infectious disease. The 1.5 million TB-associated deaths in 2020 was the first increase in over a decade^1^. Possible outcomes of pulmonary *Mycobacterium tuberculosis* (*Mtb*) infection include pathogen clearance, latent tuberculosis infection (LTBI), and active tuberculosis (ATB) disease, the latter of which results in the clinical manifestations associated with TB. Both the disease outcome and the pathology of TB are driven by the type of immune response mounted by the host. Thus, it is imperative to better understand what immune responses protect against versus promote *Mtb* infection to inform vaccine and host-directed therapy development.

Neutrophils are the most abundant and predominantly-infected cell type in the sputum^2^, bronchoalveolar lavage (BAL) fluid^3^, and caseum contents from resected lung tissue of active TB patients^2^. Studies of TB in mice^4–17^, nonhuman primates^18–21^, and humans^9, 22–28^ have identified a correlation between neutrophil abundance and increased disease severity. In contrast to their antibacterial role in numerous other infectious diseases^29^, neutrophils in TB have impaired capacity for killing phagocytosed *Mtb*^30–32^. However, there are studies supporting a protective role for neutrophils during *Mtb* infection, where neutrophils were required to restrict *Mtb* growth in human whole blood *ex vivo*^33, 34^. Isolated human neutrophils have also been shown to restrict *Mtb* replication^35^, particularly in the presence of TNFα . These data suggest that there could be a protective role for neutrophils during *Mtb* infection and highlight that the mechanisms underlying how neutrophils impact *Mtb* replication and disease progression remain open questions in the field.

In response to *Mtb* infection, neutrophils deploy a number of defenses including the release of granules containing antimicrobial molecules and the extrusion of neutrophil extracellular traps (NETs)^9, 33, 37–39^. Markers for neutrophil NETosis are present in necrotic lesions in resected lungs and in the plasma from active TB patients^9, 40–42^, suggesting an association between NETosis and active TB disease. NETosis is the process by which neutrophils undergo histone citrullination^43, 44^, chromatin decondensation^45^, and release of web-like chromatin structures decorated with antimicrobial granule proteins with the potential to bind, trap, and kill pathogens^46–48^.

*Mtb*-induced NETs are unable to kill the bacteria^38^, but can be bound and phagocytosed by macrophages to impact macrophage inflammatory responses during infection^39^.

Although very little is known about the molecular processes governing NETosis during *Mtb* infection, a recent study showed that antibody-mediated blocking of GM-CSF signaling in *Mtb*-infected mice increased neutrophil accumulation and NETosis, as indicated by staining for citrullinated histone 3 (H3Cit)^9^. Global deletion of type I interferon (IFN) receptor (IFNAR) decreased neutrophil accumulation and NETosis in GM-CSF signaling deficient mice. In addition, *Mtb* infection of C3HeB/Fej mice, which are more susceptible to infection than C57BL/6J mice and exhibit higher levels of type I IFN signaling during infection, also resulted in signs of NETosis in the lungs^9^. However, deletion of IFNAR specifically in neutrophils decreased neutrophil accumulation but not the levels of citrullinated histones in neutrophils, suggesting that type I IFN functions through other cell types to impact NETosis and leaving open the question of how type I IFN directly controls neutrophil responses. Thus, the cell biology and molecular regulation of NETosis within neutrophils during *Mtb* infection is still mostly undefined.

In this study, we dissect the cellular process of NETosis in response to *Mtb* and its direct effect on bacterial replication and pathogenesis *in vitro* and *in vivo*. We discover that during *Mtb* infection, neutrophils undergo a form of NET release where decondensed chromatin is packaged in vesicles and released from the cell in a manner that can maintain host cell viability. *Mtb*-induced NET release specifically requires peptidyl arginine deiminase 4 (PAD4)-mediated histone citrullination. Unlike PAD4, type I IFN signaling within neutrophils does not affect histone citrullination but instead promotes the formation of the chromatin containing vesicles that will be released as NETs. Analysis of markers for NETosis in *Mtb*-infected nonhuman primates (NHPs) support a model where neutrophils are recruited to necrotic regions that contain *Mtb* bacilli where they are exposed to type I IFN from epithelioid macrophages and undergo NETosis, contributing to necrosis and the formation of caseum. Furthermore, we demonstrate that NETs can directly promote *Mtb* replication and pathogenesis, thus identifying a promising pharmacological target to control both pathogen replication and the pathology associated with severe TB disease.

## RESULTS

### *Mtb* induces NET release by mouse neutrophils

To dissect the process and impact of NETosis during *Mtb* infection, we infected neutrophils isolated from wild-type (WT) C57BL/6J mice with a strain of *Mtb* Erdman expressing GFP^12, 13^ at a multiplicity of infection (MOI) of 20 and visualized NETosis using fluorescent confocal microscopy (Figure 1A). Neutrophils were identified by staining with an anti-Ly6G antibody and an antibody specific for citrullinated histone 3 (H3Cit) was used to monitor the initial step of NETosis. On average, 5.1% of neutrophils in cultures infected with *Mtb* were H3Cit positive by 4 hours post infection (hpi) as compared to 0.4% in mock infected cultures (Figures 1B and 1C). By 18 hpi with *Mtb*, 56.7% of neutrophils were H3Cit positive in contrast to 0.6% in mock infected cultures (Figures 1B and 1C). We quantified released NETs by counting extracellular H3Cit positive web-like structures using the Ridge Detection plugin^49^ of Fiji^50^ (Figure S1). We did not detect any released H3Cit positive webs in mock-infected cells at 4 hours, but 1.3 webs per 100 cells were detected at 4 hpi with *Mtb*. By 18 hpi with *Mtb*, we detected an average of 22.7 webs per 100 cells (Figure 1C). To better visualize released NET structures, we performed scanning electron microscopy (SEM) with mock or *Mtb*- infected neutrophils. We were able to detect released NET structures at both 4 and 18 hpi with *Mtb*, but not in mock infected cells at either time point (Figure 1D, red arrows). In addition, SEM revealed extracellular *Mtb* bacilli directly associated with the NETs (Figure 1D, yellow arrows). Together these data demonstrate that some murine neutrophils will release NETs in response to *Mtb* infection by 4 hpi, and this number of NETotic neutrophils increases in response to *Mtb* by 18 hpi. These results also provide the first evidence of *Mtb* induced NETosis by mouse bone marrow neutrophils *in vitro*.

**Figure 1.**
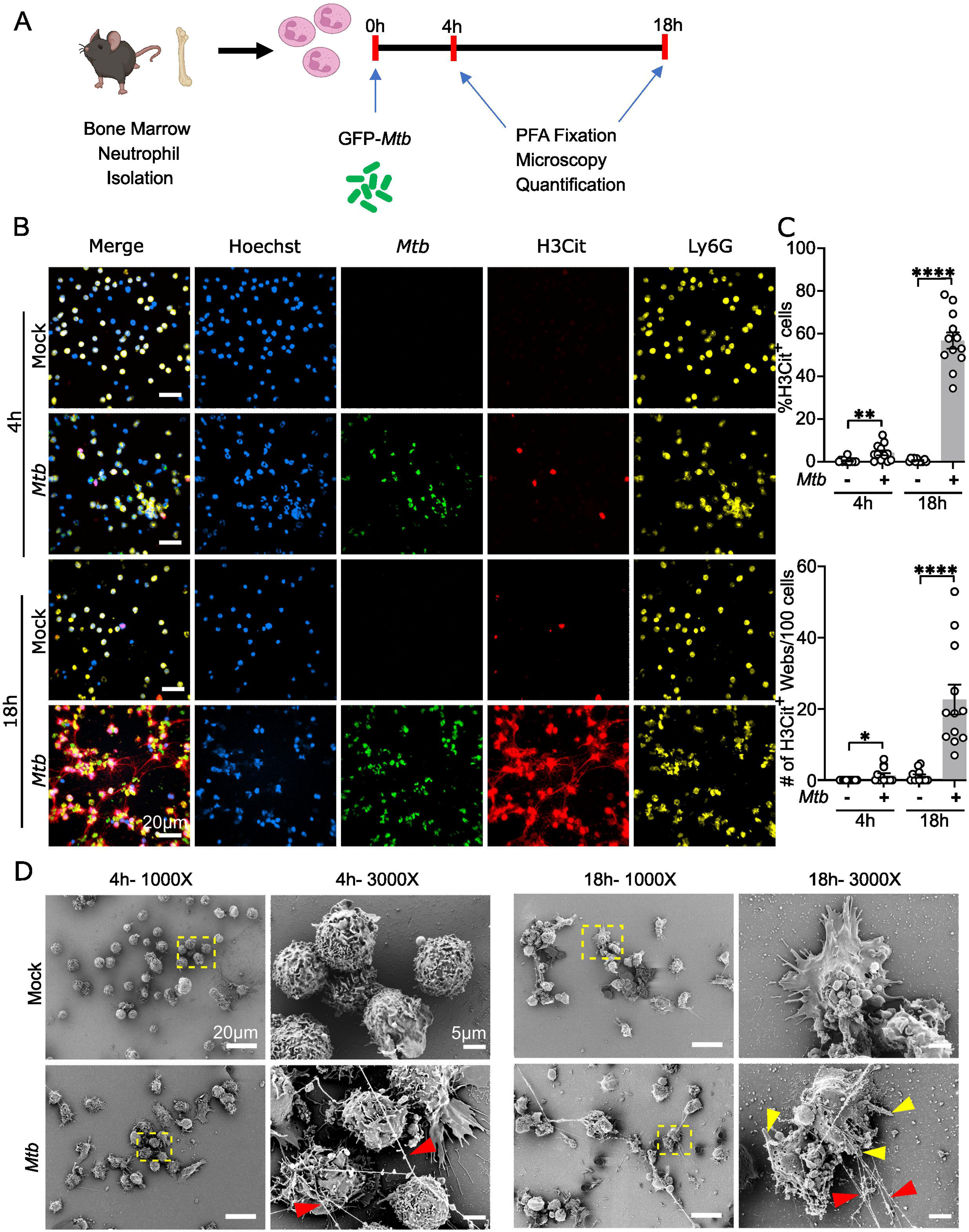
*Mtb* induces NET formation by mouse neutrophils. **(A)** Schematic of experimental design for *in vitro Mtb* infection of murine bone marrow neutrophils. Bone marrow neutrophils from C57BL/6J mice were cocultured with normal mouse serum (NMS)-opsonized GFP-*Mtb* at an MOI of 20 or NMS (mock) for 4h and 18h before fixing with PFA and visualizing by confocal microscopy. BioRender.com used in schematic. **(B)** Representative confocal images of neutrophils stained for citrullinated histone 3 (H3Cit), the neutrophil marker Ly6G, and DNA (Hoechst). GFP-*Mtb* is also shown. Magnification 60x. **(C)** The percentage of Hoechst^+^ cells that were also H3Cit^+^ (top) and the number of extracellular H3Cit^+^ webs per 100 Hoechst^+^ nuclei (bottom) per field under 60x objective were quantified using ImageJ software and plotted. Each datapoint represents data from one field and a total of 12 fields containing 20-200 cells/field were quantified and compiled from 4 independent experiments. Bar graph of data represents mean ± SEM. ****P < 0.0001 determined by unpaired t test, comparing only within a L single timepoint. **(D)** Representative scanning electron microscopy showing neutrophil morphology after 4 and 18 hpi with *Mtb* or following mock infection. Released NETs (red arrows) are observed in association with extracellular *Mtb* (yellow arrows). Images on the right for each time point are zoomed in from the region in the yellow box.

### NETs released in response to *Mtb* infection differ structurally from PMA and ionomycin induced NETs

To determine whether NETosis in response to *Mtb* infection resembled NETosis in response to other stimuli, we compared neutrophils infected with *Mtb* to neutrophils treated with two well-studied chemical stimuli of NETosis, phorbol myristate acetate (PMA) and ionomycin. PMA activates phosphokinase C (PKC) and ERK signaling, which induces NADPH oxidase-dependent NETosis^46, 51^. In contrast, ionomycin increases cytosolic calcium levels, which induces NETosis independent of NADPH oxidase^52, 53^. Compared to PMA and ionomycin, *Mtb* induced significantly less histone citrullination after 4 hours of incubation (Figure S2A and S2B), however, after 18 hours of infection or treatment, all three conditions induced >50% H3Cit positivity, with *Mtb* infection resulting in a significantly higher percentage of H3Cit positive cells compared to PMA (Figures 2A and 2B). Despite the higher percentage of H3Cit positive cells, *Mtb* infection and PMA treatment resulted in a similar number of released NETs. The most striking observation was the difference in the thickness of the NETs released, where *Mtb* infection led to the release of significantly thinner strands of H3Cit coated DNA than PMA or ionomycin treatment (Figures 2A and 2B). SEM imaging of *Mtb*-infected and PMA-treated neutrophils provided a higher resolution of the NET ultrastructure and revealed that neutrophils released thin threads of DNA in response to *Mtb* infection whereas PMA treatment resulted in the release of larger chromatin bundles (Figure 2C). In addition, contrary to PMA treatment that induced cell flattening of neutrophils, *Mtb* infected neutrophils retained their round morphology during the process of NETosis (Figures 2C and S2C), suggesting different mechanisms of NETosis in responses to these two stimuli.

**Figure 2.**
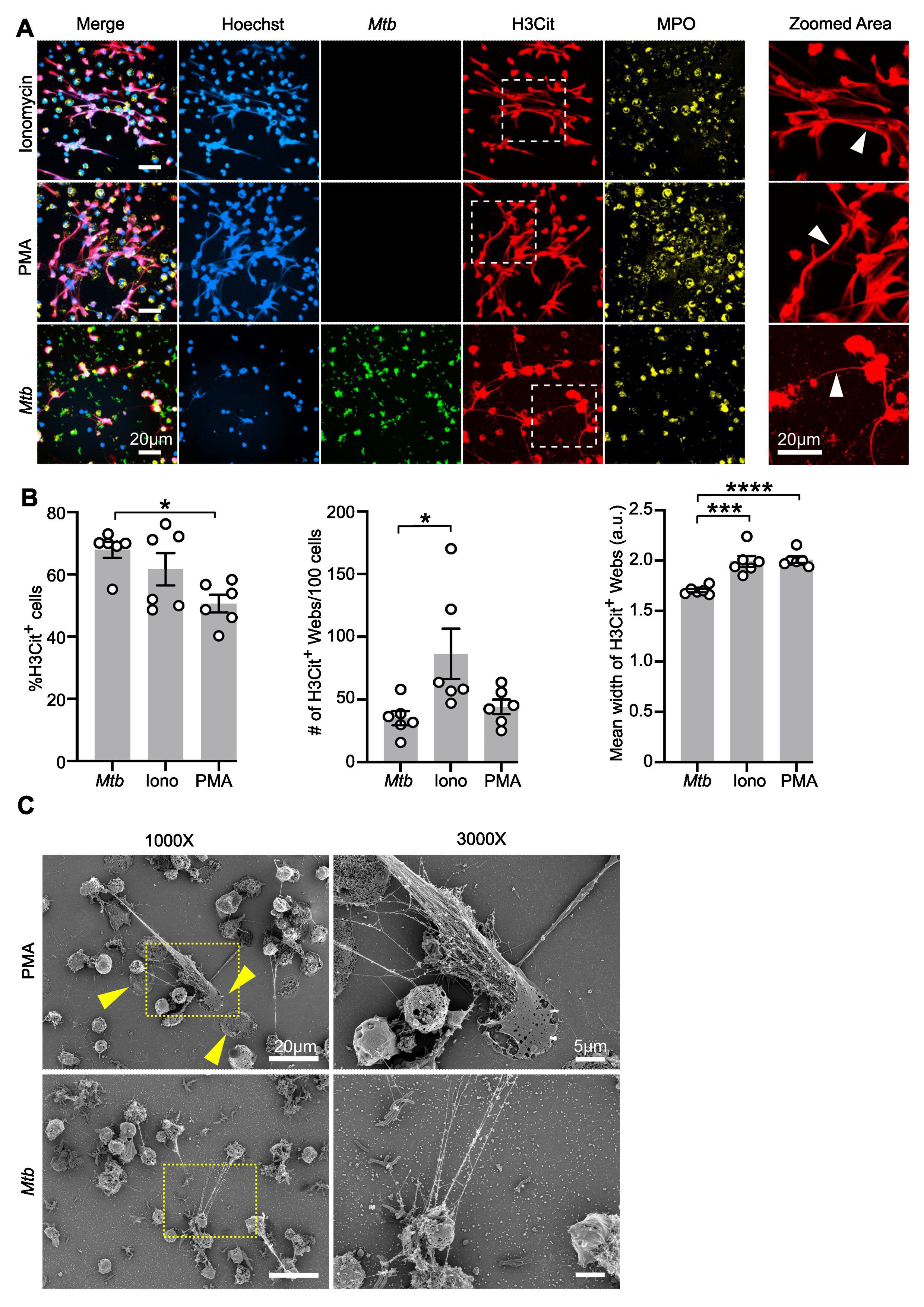
NETs released in response to *Mtb* infection differ structurally from PMA and ionomycin induced NETs. **(A)** Representative confocal images showing immunofluorescence staining of PFA fixed bone marrow neutrophils from C57BL/6J mice infected with *Mtb* at an MOI 20 or treated with PMA (100 nM) or ionomycin (5 µM) for 18 h and stained for citrullinated histone 3 (H3Cit), the neutrophil marker myeloperoxidase (MPO), and DNA (Hoechst). GFP-*Mtb* is also shown. Images on the right are zoomed in from the region in the white box. White arrows denote NETs formed in each condition. Magnification 60x, zoomed in 2x. **(B)** The percentage of Hoechst^+^ cells that were also H3Cit^+^ (left), the number of extracellular H3Cit^+^ webs per 100 Hoechst^+^ nuclei (middle), and mean width of H3cit^+^ webs (right) per field at 18 h of *Mtb* infection or treatment with PMA or ionomycin (Iono) under 60x objective were quantified using ImageJ software and plotted. Each datapoint represents data from one field and a minimum of 6 fields containing 20-200 cells/area were quantified and compiled from 2 independent experiments. Bar graph of data reprsents mean ± SEM. *P < 0.05; ***P < 0.001; ****P < 0.0001 by one-way ANOVA with Tukey’s correction. **(C)** Representative scanning electron microscopy showing neutrophil morphology after 18 h of coculture with *Mtb* or treatment with PMA. Yellow arrows denote the flattening of neutrophils in the presence of PMA. Clustered release of chromatin was observed during PMA treatment compared to thread like NETs released during *Mtb* infection. Images on the right are zoomed in from the region in the yellow box.

### *Mtb* infected neutrophils package chromatin in vesicles for release and can maintain viability during NET release

NETosis has historically been defined as a suicidal cell death process^47^. However, the thinner NETs released from *Mtb* infected neutrophils and the observation that the *Mtb*- infected NETotic neutrophils maintain their round morphology (Figures 1D and 2C) is reminiscent of the recently described process of vital NET release where the NETotic neutrophil retains its viability along with effector functions^54, 55^. To investigate if neutrophils maintain viability during NET release in response to *Mtb* infection, we included the Zombie Viability Dye (BioLegend) to stain cells with compromised plasma membrane integrity in our microscopy experiments. At 18 hpi with *Mtb* there was a significantly lower frequency of (21%) Zombie Dye positive cells than observed in cultures following 18 hours of PMA or ionomycin treatment (69% and 84% Zombie^+^, respectively), despite having similarly high levels of H3Cit positivity and released NETs (Figures 3A and 3B). When we specifically quantified the viability of H3Cit^+^ neutrophils, only an average 18% of H3Cit^+^ cells were Zombie^+^ at 18 hpi with *Mtb*, as compared to an average of over 90% H3Cit^+^ cells being Zombie^+^ after PMA or ionomycin treatment (Figure 3B). These data indicate that some neutrophils maintain viability following NET release in response to *Mtb* infection.

**Figure 3.**
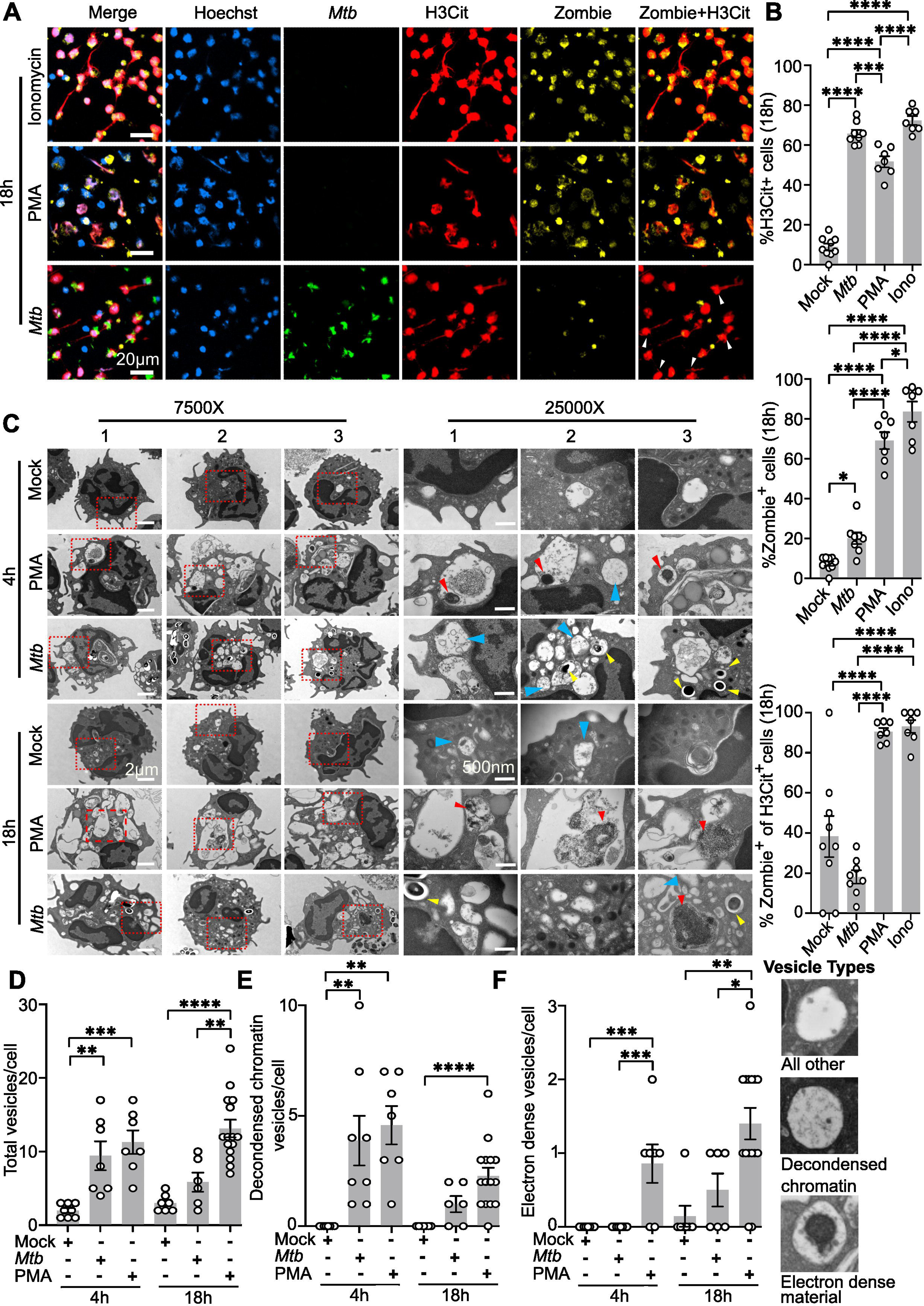
*Mtb* infected neutrophils package chromatin in vesicles for release and can maintain viability during NET release. **(A)** Representative confocal images showing bone marrow neutrophils from C57BL/6J mice infected with GFP-*Mtb* at an MOI 20 or treated with PMA (100nM) or ionomycin (Iono, 5µM) for 18h and stained for citrullinated histone 3 (H3Cit), cell death marker (Zombie dye), and DNA (Hoechst). GFP-*Mtb* is also shown. White arrows indicate Zombie negative neutrophils positive for H3Cit staining and associated with released NETs. Magnification 60x, zoomed in 2x. **(B)** Bar graphs showing quantification of immunofluorescence staining to detect H3Cit positivity and neutrophil viability. The percentage of Hoechst^+^ cells that were H3Cit^+^ (top), the percentage of Hoechst^+^ cells that were Zombie^+^ (middle), and the percentage of H3cit^+^ cells that were Zombie^+^ (bottom) per field under 60x objective were quantified using ImageJ software and plotted. Each datapoint represents data from one field and a minimum of 6 fields containing 30-200 cells/field for each condition were quantified and compiled from 2 independent experiments. Bar graph of data represents < 0.0001 by one-way ANOVA with Tukey’s correction. < 0.05; ***P < 0.001; ****P< 0.0001 determined L by one-way ANOVA with Tukey’s correction. **(C)** Representative TEM images of neutrophils infected with *Mtb* at an MOI of 20, mock- infected, or treated with PMA for 4h or 18h. Three representative cells per condition are shown. Red boxes designate the region that was zoomed into for each cell and shown in the right panels. Yellow arrows denote *Mtb*, blue arrows denote vesicles containing “beads on a string” DNA and histone structures (decondensed chromatin vesicles), and red arrows denote vesicles containing electron dense material. **(D-F)** Quantification of different types of vesicles observed in neutrophils using TEM after PMA treatment, *Mtb* infection, or mock infection. (D) Total vesicles per cell, (E) vesicles containing decondensed chromatin, and (F) vesicles containing electron dense material per cell at 4h and 18h following treatment or infection. Each datapoint represents a single neutrophil and 6-20 neutrophils were used for quantification per sample group from two independent experiments. Bar graph of data represents mean ± SEM. *P < 0.05; ***P < 0.001; ****P < 0.0001 by one-way ANOVA with Tukey’s correction, comparing only within a given timepoint.

Vital NET release occurs via nuclear envelope blebbing of decondensed chromatin into vesicles that are subsequently exocytosed from the cell^54, 56, 57^. To determine if *Mtb*- induced NETosis exhibits similar subcellular morphological features to vital NET release, we performed ultrastructure analysis of neutrophils after 4 and 18 hours of infection with *Mtb*, mock infection, or treatment with PMA using transmission electron microscopy (TEM) (Figure 3C). *Mtb* infection and PMA treatment induced increased vesicle formation by 4 hours as compared to mock infected controls (Figures 3C and 3D). By 4 hours we were also able to identify vesicles containing decondensed chromatin consisting of DNA strands exhibiting a “beads on a string” appearance in both *Mtb*-infected and PMA-treated cultures, but not in mock infected cultures (Figures 3C (blue arrows), 3E, and S3A). The number of vesicles containing decondensed chromatin per cell was similar in cultures following *Mtb* infection or PMA treatment at both 4 hours and 18 hours (Figure 3E), indicating that this was not a feature unique to maintained viability during NET release. Some chromatin containing vesicles also contained *Mtb* (Figure 3C), suggesting that vesicles containing chromatin were fusing with *Mtb*-containing vesicles. During both *Mtb* infection and PMA treatment, release of vesicles through the plasma membrane could be observed (Figures S3B,C). The only morphological feature noted to be different during PMA treatment versus *Mtb* infection was that by 4 hours the PMA-treated cells contained vesicles harboring electron dense material that were not observed in *Mtb* infected cells at this time point (Figures 3C and 3F). By 18 hpi with *Mtb*, some cells had formed the vesicles containing electron dense material, but still to a lesser extent than during PMA treatment at this time point (Figure 3F). Therefore, the abundance of the electron dense material is correlated with increased cell death during NETosis. We were able to identify some PMA-treated cells with vesicles that contained the electron dense material uncoiling into decondensed “beads on a string” chromatin (Figure S3D), suggesting that this electron dense material could represent condensed chromatin. Together, these results support a model where during *Mtb* infection, neutrophils release NETs via vesicles in a manner that can preserve plasma membrane integrity.

### Histone citrullination during *Mtb*-induced NET release is PAD4-dependent

Histone citrullination by PAD enzymes is critical for the initial chromatin decondensation that allows for NETosis following most stimuli^58–60^, including during *Mtb* infection where pretreatment with the pan-PAD inhibitor Cl-amidine inhibited *Mtb* induced NETosis by human neutrophils^61^. PAD4 is the primary PAD enzyme expressed in neutrophils and has been shown to be essential for NETosis in response to a number of different stimuli^44, 52, 59, 60, 62, 63^ including lipopolysaccharide, lipoteichoic acid, fungal zymosan, and TNFα However, NET formation in response to other stimuli, such as *Klebsiella pneumoniae, Aspergillus fumigatus,* rodent-specific pneumovirus, and influenza virus A/WSN/33/H1N1, occurs independent of PAD4^64–68^. To determine if *Padi4*, the gene that encodes PAD4, is required for NETosis during *Mtb* infection, we infected WT and *Padi4*^-/-^ neutrophils with *Mtb* for 18 hours and monitored NETosis by fluorescent microscopy. Genetic inhibition of *Padi4* led to an almost complete loss of H3Cit positivity and released NETs during *Mtb* infection (Figures 4A and 4B). We also quantified H3Cit positivity by flow cytometry and observed a significantly lower frequency of H3Cit^+^ *Padi4*^-/-^ neutrophils compared to WT neutrophils at 18 hpi (Figure 4C). Moreover, PAD4-deficient neutrophils harbored a significantly reduced number of DNA-filled vesicles per cell upon stimulation with *Mtb* at 18 hpi (Figures 4D-E), demonstrating that PAD4 is required for histone citrullination and NETosis during *Mtb* infection.

**Figure 4.**
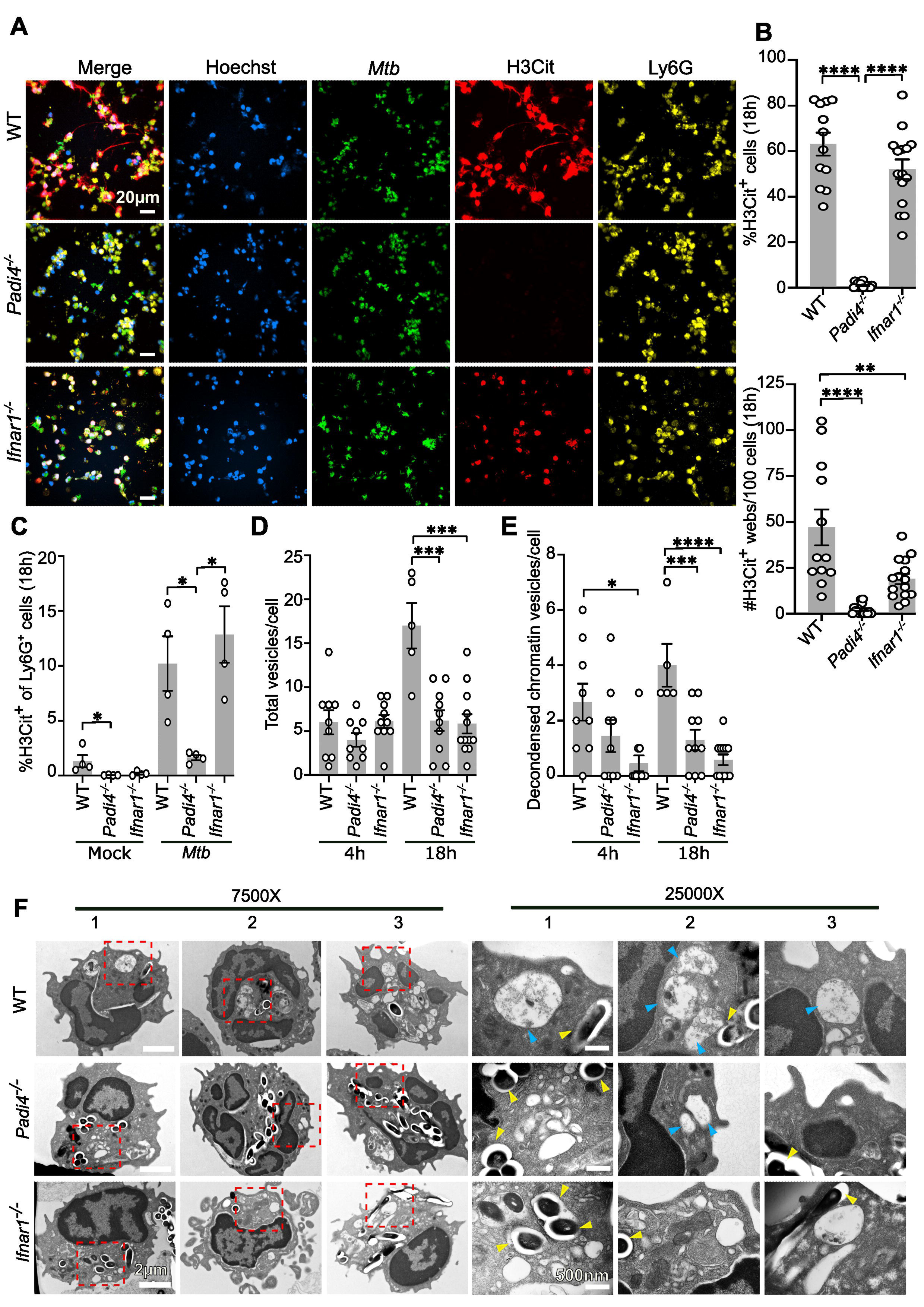
Molecular determinants of NETosis during *Mtb* infection. **(A)** Representative confocal images showing immunofluorescence staining of neutrophils from WT, *Padi4*^-/-^, and *Ifnar1*^-/-^ mice infected with GFP-*Mtb* at an MOI of 20 for 18h and stained for citrullinated histone 3 (H3Cit), a neutrophil marker (Ly6G), and DNA (Hoechst). GFP-*Mtb* is also shown. Magnification 60x. **(B)** The percentage of Hoechst^+^ cells that were also H3Cit^+^ (top) and the number of extracellular H3Cit^+^ webs per 100 Hoechst^+^ nuclei (bottom) per field under 60x objective were quantified using ImageJ software and plotted. Each datapoint represents a single field and a minimum of 12 fields containing 20-200 cells/field were quantified and compiled from 2 independent experiments. Bar graph of data represents the mean ± SEM. *P < 0.05; ***P < 0.001; ****P < 0.0001 by one-way ANOVA with Tukey’s correction. **(C)** Bar graph represents the percent of Ly6G^+^ cells that were also H3Cit^+^ as determined by flow cytometry after 18 hpi with *Mtb* or mock infected. Neutrophils were collected from WT, *Padi4*^-/-^, and *Ifnar1*^-/-^ mice where each datapoint for a given strain and condition represents a different mouse. N=4 mice for each genotype. Bar graph of data represents mean ± SEM. *P < 0.05 by one-way ANOVA with Tukey’s correction, compared within a single condition. **(D-E)** Bar graphs showing quantification of different types of vesicles observed in neutrophils using TEM after *Mtb* infection. (D) Total vesicles per cell and (E) vesicles containing decondensed chromatin at 4h and 18h following infection. Each datapoint represents a single neutrophil, where 5-10 neutrophils were used for quantification per L mean ± SEM. *P < 0.05; ***P < 0.001; determined by one-way ANOVA with Tukey’s correction, comparing < 0. only within a given timepoint. **(F)** Representative TEM images of bone marrow neutrophils from WT, *Padi4*^-/-^ and *Ifnar1*^-/-^ mice infected with *Mtb* for 4h. Three representative cells per strain are shown. Red boxes designate the region that was zoomed into for each cell and shown in the right panels. *Mtb* (yellow arrow) and vesicles containing decondensed chromatin (blue arrow) are labeled.

### Type I IFN regulates NET release, but not histone citrullination, during *Mtb* infection

Type I IFN signaling was shown to impact levels of NETosis in susceptible mice, but this role was proposed to not be neutrophil intrinsic based on retained H3Cit staining in mice deleted for *Ifnar1* specifically in neutrophils^9^. Type I IFN signaling is associated with increased NETosis in multiple other contexts as well^69, 70^, although the exact role for type I IFN signaling is still unknown in all cases. We directly investigated whether type I IFN signaling within neutrophils impacts NETosis during *Mtb* infection by infecting WT and *Ifnar1*^-/-^ neutrophils with *Mtb* for 18 hours and monitoring histone citrullination and NET release by microscopy and flow cytometry. Similar to what was observed *in vivo*^9^, loss of type I IFN signaling in neutrophils had no effect on the levels of H3Cit in neutrophils during *Mtb* infection (Figures 4A-C). However, we observed a significant decrease in the released NETs from *Ifnar1*^-/-^ neutrophils compared to WT neutrophils during *Mtb* infection (Figures 4A and 4B). These data suggested that type I IFN signaling in neutrophils regulates a step of NETosis after histone citrullination. Therefore, we investigated whether loss of type I IFN signaling was affecting the formation of vesicles containing decondensed chromatin. *Mtb* infection of *Ifnar1*^-/-^ neutrophils resulted in significantly fewer chromatin filled vesicles per cell by 4 hpi as compared to WT neutrophils (Figures 4D-F). At 18 hpi, the numbers of total vesicles and vesicles containing decondensed chromatin per cell was still lower in *Ifnar1*^-/-^ neutrophils compared to WT neutrophils, indicating that the defect in vesicle number was not merely a delay in the NETosis process, but was a block prior to vesicle formation. Together these studies demonstrate that histone citrullination during *Mtb* infection is PAD4-dependent, whereas the release of citrullinated DNA is regulated by type I IFN signaling in neutrophils. This newly discovered neutrophil-intrinsic role for type I IFN in NET release could also explain why previous studies observed that deletion of *Ifnar1* in neutrophils resulted in decreased tissue pathology without affecting H3Cit levels^9^.

### NETs directly promote replication of *Mtb in vitro* and *in vivo*

Depletion of neutrophils in some susceptible mice can reverse high bacterial burdens^8, 9, 11, 12^, however, it remains unknown how mechanistically neutrophils elicit effects on *Mtb* replication. We examined whether NETs could directly contribute to *Mtb* survival and replication by infecting WT and *Padi4*^-/-^ neutrophils with *Mtb* and monitoring bacterial burden at 48 hpi by plating for colony forming units (CFU). *Mtb* burdens were higher in cultures containing neutrophils than when *Mtb* was grown alone in the same media (Figure 5A), indicating that neutrophils can directly promote *Mtb* replication. In contrast, the higher bacterial burdens were completely reversed in cultures of *Mtb*- infected *Padi4*^-/-^ neutrophils, where *Mtb* grew to similar levels as in cultures lacking neutrophils (Figure 5A). These data indicate that NETosis is a contributor to *Mtb* replication in the presence of neutrophils. In addition, deletion of *Ifnar1* also resulted in lower levels of *Mtb* replication compared to WT neutrophils (Figure 5A), supporting a role for NETosis in promoting *Mtb* replication.

**Figure 5.**
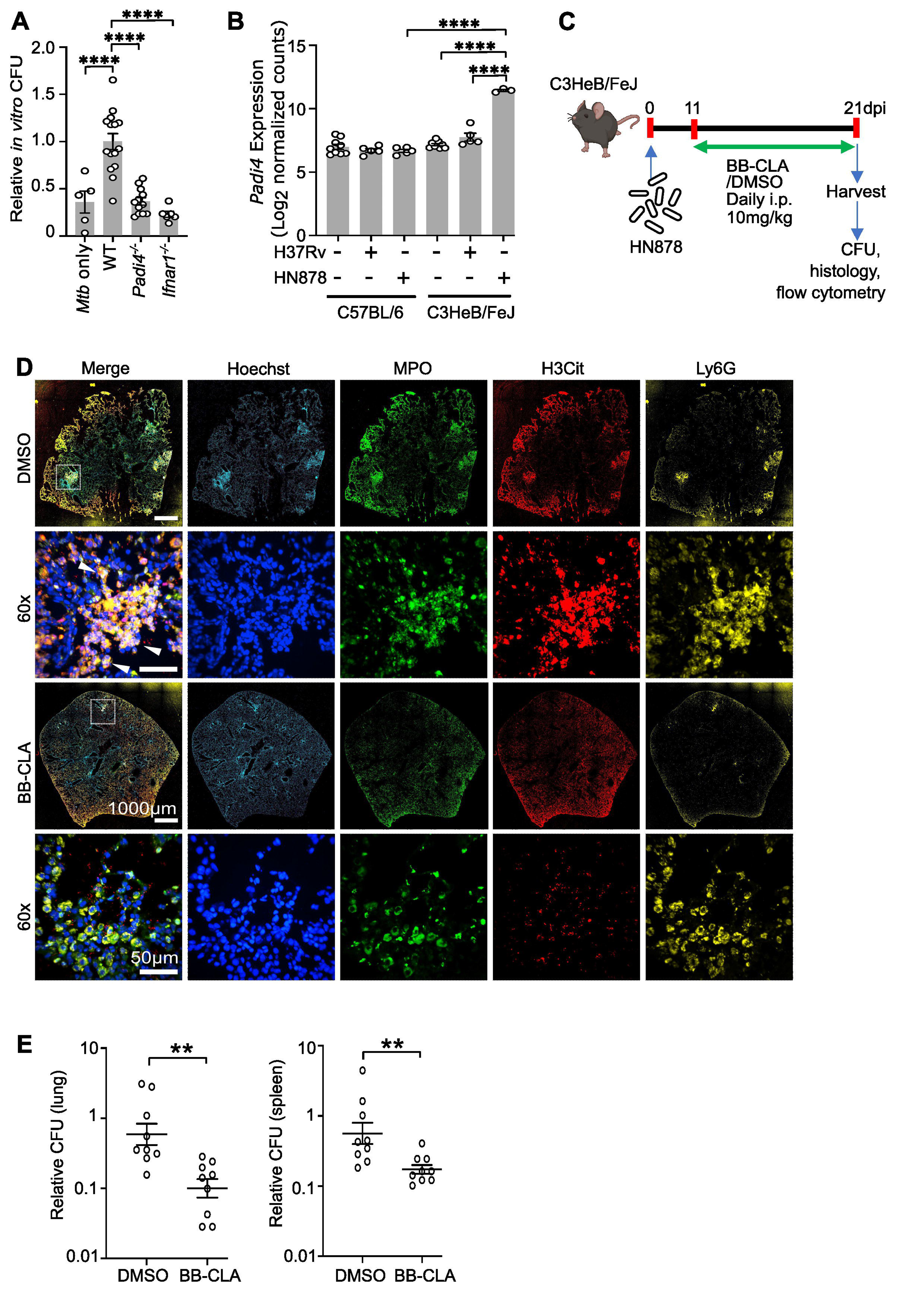
NETs directly promote replication of *Mtb in vitro* and *in vivo*. **(A)** Bone marrow neutrophils from WT, *Padi4*^-/-^, and *Ifnar1*^-/-^ mice were infected with GFP-*Mtb* at an MOI of 20 or an equivalent amount of *Mtb* was cultured in the same media without neutrophils for 48 hours. CFU/ml were determined by lysing the neutrophils and plating dilutions of the entire well on 7H11 plates at 48 hours. Each datapoint in the graph is the CFU/ml for each sample expressed relative to the average CFU/ml in *Mtb*-infected WT neutrophil cultures within the same experiment. Each datapoint is from a single well compiled from two independent experiments. Bar graph of data represents mean ± < 0.0001 by one-way ANOVA with Tukey’s correction. BioRender.com used in schematic. **(B)** Analysis of normalized gene expression data from lungs of C57BL/6J or C3HeB/FeJ mice uninfected or infected with low dose Lineage 4 H37Rv *Mtb* strain or Lineage 2 HN878 *Mtb* strain. The expression data was obtained from https://ogarra.shinyapps.io/tbtranscriptome/. Bar graph of data represents mean, where *****P < 0.0001 by one-way ANOVA with Tukey’s correction. **(C)** Schematic of experimental design for *Mtb* HN878 aerosol infection of C3HeB/FeJ mice followed by intraperitoneal injection with 10 mg/kg of BB-Cl-amidine or DMSO vehicle every other day from day 11 post-infection until lungs and spleens were harvested at 21 dpi for analysis. N=9 mice per condition compiled from 2 independent experiments. **(D)** Representative images showing immunofluorescence staining of lung sections from *Mtb*-infected C3HeB/FeJ mice that were probed with antibodies to detect citrullinated histone H3 (H3Cit; red), myeloperoxidase (MPO, green, a marker for neutrophil granules), Ly6G (yellow, a marker for neutrophils) and DNA (Hoechst, blue). The entire lung section is shown along with a 60x zoomed in region denoted by the white box. The white arrows point to colocalization of H3Cit, DAPI, and Ly6G staining, indicating the presence of NETing neutrophils. **(E)** Bacterial burdens in the lungs (left) and spleen (right) at 21 dpi were determined by plating dilutions of the organ homogenate onto 7H11 agar plates to numerate CFUs. Each datapoint is the CFU/ml for a single mouse relative to the average CFU/ml in DMSO vehicle treated mice within the same experiment. Data are log_10_ transformed and the error bars represent the mean ± SEM. **P 01 determined by unpaired t test.

Publicly available transcriptomic data from the lungs of *Mtb*-infected mice^71^ indicates that *Mtb* infection does not upregulate *Padi4* expression in WT C57BL/6J *in vivo* (Figure 5B). In contrast, C3HeB/FeJ mice infected with the Lineage 2 *Mtb* strain HN878 significantly induced *Padi4* transcript production in their lungs compared to uninfected mice or infected C57BL/6J mice (Figure 5B)^71^. H3Cit signal has also been detected in the lungs of *Mtb*-infected C3HeB/FeJ mice in prior studies^9^. Therefore, to investigate how NETosis impacts *Mtb* replication *in vivo*, we infected C3HeB/FeJ mice with *Mtb* HN878 and chemically inhibited NETosis with daily intraperitoneal (IP) injections with the pan-PAD inhibitor BB-Cl-amidine (Cayman Chemical Company) (Figures S4A and S4B) starting at 11 dpi until harvesting lungs for analysis of inflammation, histone citrullination, and bacterial burdens at 21 dpi (Figure 5C). Histological analysis of lung lesions in BB-Cl-amidine or DMSO vehicle-treated *Mtb*-infected mice demonstrated that neutrophils accumulated in the lungs of all infected mice, but there was significantly more H3Cit staining in the DMSO vehicle-treated *Mtb*-infected mice (Figures 5D and S5), indicating that BB-Cl-amidine was effective at blocking histone citrullination in neutrophils during *Mtb* infection of C3HeB/FeJ mice. Flow cytometry analysis of lungs at 21 dpi revealed that BB-Cl-amidine-treated *Mtb*-infected C3HeB/FeJ mice accumulated more neutrophils and B cells in the lungs, but did not exhibit any other significant differences in cell populations as compared to DMSO vehicle-treated *Mtb*-infected mice (Figure S4C). Blocking NETosis in *Mtb*-infected C3HeB/FeJ mice with BB-Cl-amidine resulted in significantly lower bacterial burdens in the lungs and spleens at 21 dpi (Figure 5E), despite still accumulating high levels of neutrophils (Figure S4C), suggesting that NETosis directly contributes to *Mtb* replication *in vivo* and can be chemically inhibited to control *Mtb* pathogenesis.

### NETosis is associated with necrotic microenvironments in primate granulomas

In addition to the effects of NETosis on *Mtb* replication, we were interested in how NETosis contributes to granuloma-level pathology in humans. While most mouse strains do not develop the diverse range of lesion types observed in humans, *Mtb*-infected cynomolgus macaques experience the full spectrum of pathology seen in human TB and develop granulomas that are equivalent to their human counterparts^19, 72^. Macaque granulomas contain all the microenvironments that human granulomas do, including non-diseased lung adjacent to the granuloma, the T- and B cell-rich lymphocyte cuff, the epithelioid macrophage region, and, in many granulomas, a necrotic core^73^. Neutrophils experience different stimuli in each region^21, 72^ and we took advantage of this feature in granulomas from cynomolgus macaque with active TB to identify relationships between NETosis and different granuloma microenvironments.

We found substantial variation in the abundance of H3Cit^+^ and H3Cit^-^ neutrophils per granuloma (Figure 6A-C) in lesions from cynomolgus macaques. H3Cit^+^ neutrophils were abundant at the interface of epithelioid macrophages and caseum in necrotic granulomas (Figure 6B) whereas non-necrotic granulomas contained far fewer H3Cit^+^ cells (Figure 6C). When we quantified the number of H3Cit^+^ cells/granuloma, we found that non-necrotic granulomas contained significantly fewer H3Cit^+^ cells than necrotic granulomas (median ± SEM: non-necrotic granulomas 0.0136±0.0036 versus necrotic granulomas 0.0863±0.0160; Figure S6). Granulomas in close spatial proximity often had substantially different phenotypes regarding the abundance of NETotic neutrophils (Figure 6A) and we noted these differences could occur within the same granuloma if that granuloma contained multiple neutrophilic or necrotic foci (Figure 6D).

**Figure 6.**
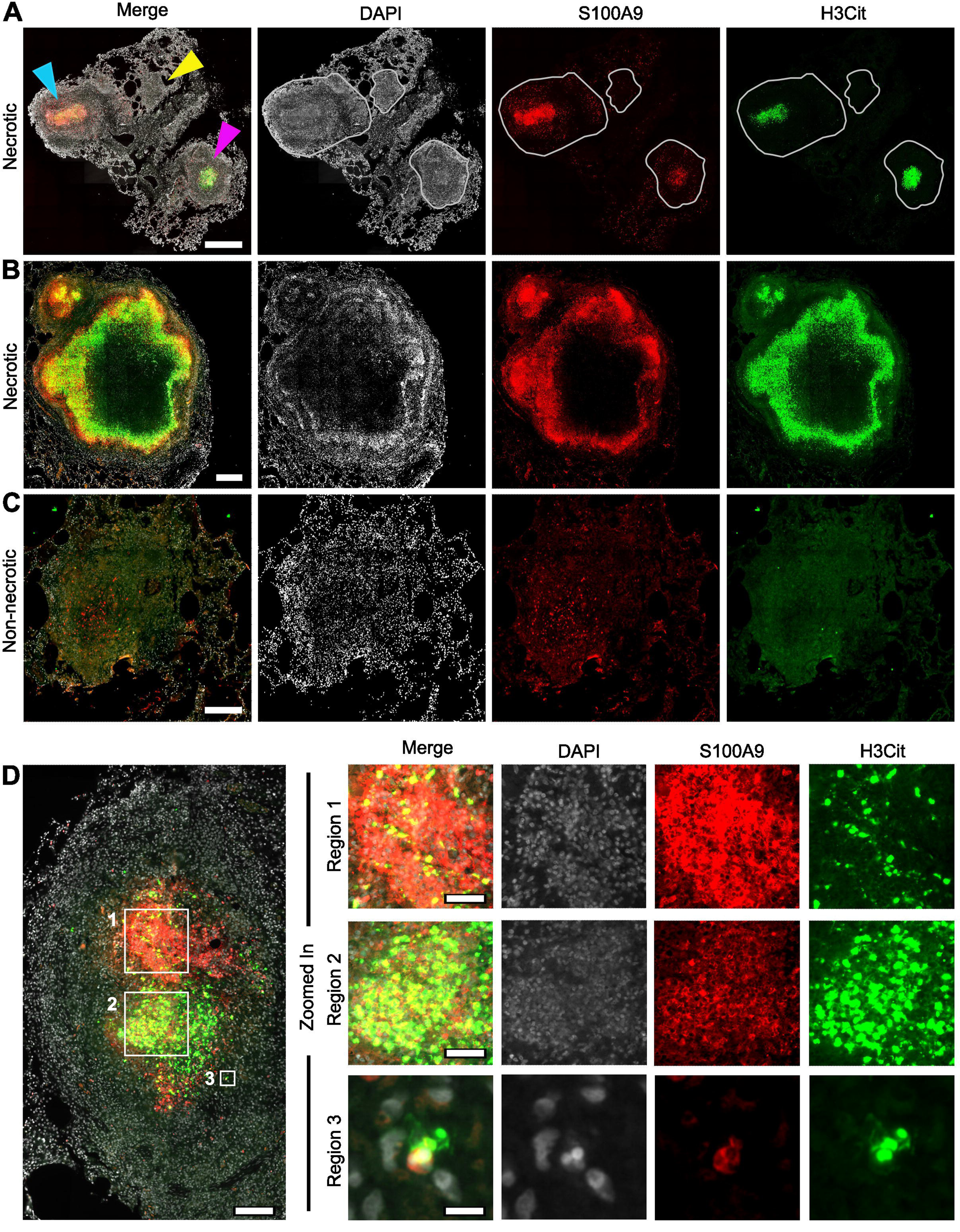
NETosis occurs in necrotic granulomas from *Mtb*-infected cynomolgus macaques. **(A)** Lung tissue from a cynomolgus macaque containing two necrotic granulomas (blue and pink arrowheads) and a non-necrotic granuloma (yellow arrowhead) was stained to visualize neutrophils (S100A9; red), H3Cit (green), and nuclei (DAPI, grey). Few H3Cit^+^ cells and neutrophils are present in the non-necrotic granuloma, whereas the necrotic granulomas are characterized by their robust populations of H3Cit^+^ neutrophils. 20x magnification; scale bar represents 500 μm. **(B)** A necrotic granuloma (larger lesion, center) with a small necrotizing lesion (upper left) that is disseminated from the larger lesion where each granuloma contains numerous H3Cit^+^ and H3Cit^-^ neutrophils in regions adjacent to caseum. 20x magnification, scale bar represents 500 μm. **(C)** H3Cit^+^ neutrophils are largely absent from a non-necrotic granuloma. 20X magnification; scale bar represents 250 μm. **(D)** Adjacent foci in a neutrophilic granuloma stained for nuclei (DAPI, grey), S100A9 (red), and H3Cit (green) (left panel) suggest that neutrophil recruitment and NETosis are dynamic processes. Region 1 (top panels, right) indicates a neutrophilic region with intact-appearing nuclei, strong S100A9 staining, and sparse H3Cit staining that contrasts with Region 2 (middle panels, right), a necrotic region with diffusely stained DNA suggestive of nuclear breakdown, dim S100A9 staining, and strong H3Cit expression. Scale bar represents 100 μm or 40 μm for the left and right panels, respectively. Region 3 highlights a S100A9^+^ neutrophil that is releasing web-like H3Cit^+^ DNA. Region 3 scale bar represents 10 μm.

We also noted an inverse relationship between S100A9 (our neutrophil marker) and H3Cit expression. This was seen in regions where strong S100A9 staining coincided with intact-appearing nuclei and less H3Cit staining (Figure 6D, region 1) whereas neutrophils in regions with less S100A9 staining had DNA that was more diffuse and stronger H3Cit expression (Figure 6D, region 2). This inverse relationship is also visible when independent necrotic granulomas were compared (Figure 6A; blue arrowhead: H3Cit^low^S100A9^high^, pink arrowhead: H3Cit^high^S100A9^low^). When we examined individual neutrophils undergoing NETosis in tissue adjacent to granuloma’s central regions (Figure 6D, region 3), we observed that H3Cit^+^ DNA appeared as if it was being discharged from these cells and this was accompanied by a flow of S100A9 protein in the same direction as the H3Cit^+^ DNA, suggesting that cytoplasmic contents were being released into the extracellular milieu as part of this process *in vivo*. Taken together, our observations demonstrate that NETosis is a terminal event for neutrophils recruited to necrotic primate granulomas but also highlight that this outcome varies by granuloma and is related to the granuloma’s morphology. Moreover, NETotic neutrophils are important contributors to the milieu of highly-degraded DNA in caseum, thus providing a link between NETosis and caseation in necrotic granulomas.

### NETosis occurs in proximity to caseum and IFN**α**2-expressing epithelioid macrophages

Our *in vitro* data demonstrated that type I IFN induces NET release during *Mtb* infection. Therefore, we investigated how NETosis within granulomas relates to type 1 IFN expression by staining macaque granulomas for IFNα2, H3Cit, and CD11c, an antigen that is broadly expressed by granuloma macrophages (Figure 7). As previously noted, H3Cit^+^ cells were present in the space between caseum and CD11c+ epithelioid macrophages (Figure 7A, left). When we visualized the IFNα2 fluorescence, we found that many epithelioid macrophages stained positively for IFNα2 (Figure 7A, right). To confirm the likelihood that these cells were producing IFNα2, we also stained these lesions for phosphorylated IRF3 (pIRF3), a transcription factor that regulates IFNα2 expression^74^. The pIRF3 staining mirrored our IFNα2 staining in epithelioid macrophages (Figure 7B, right), indicating that the epithelioid macrophages surrounding the H3Cit^+^ cells are expressing IFNα2. Furthermore, we confirmed that although neutrophils may be present throughout a granuloma (Figure 7B), H3Cit^+^ neutrophils were restricted to necrotic regions (Figure 7A) between the IFNα2^+^ epithelioid macrophages and caseum.

**Figure 7.**
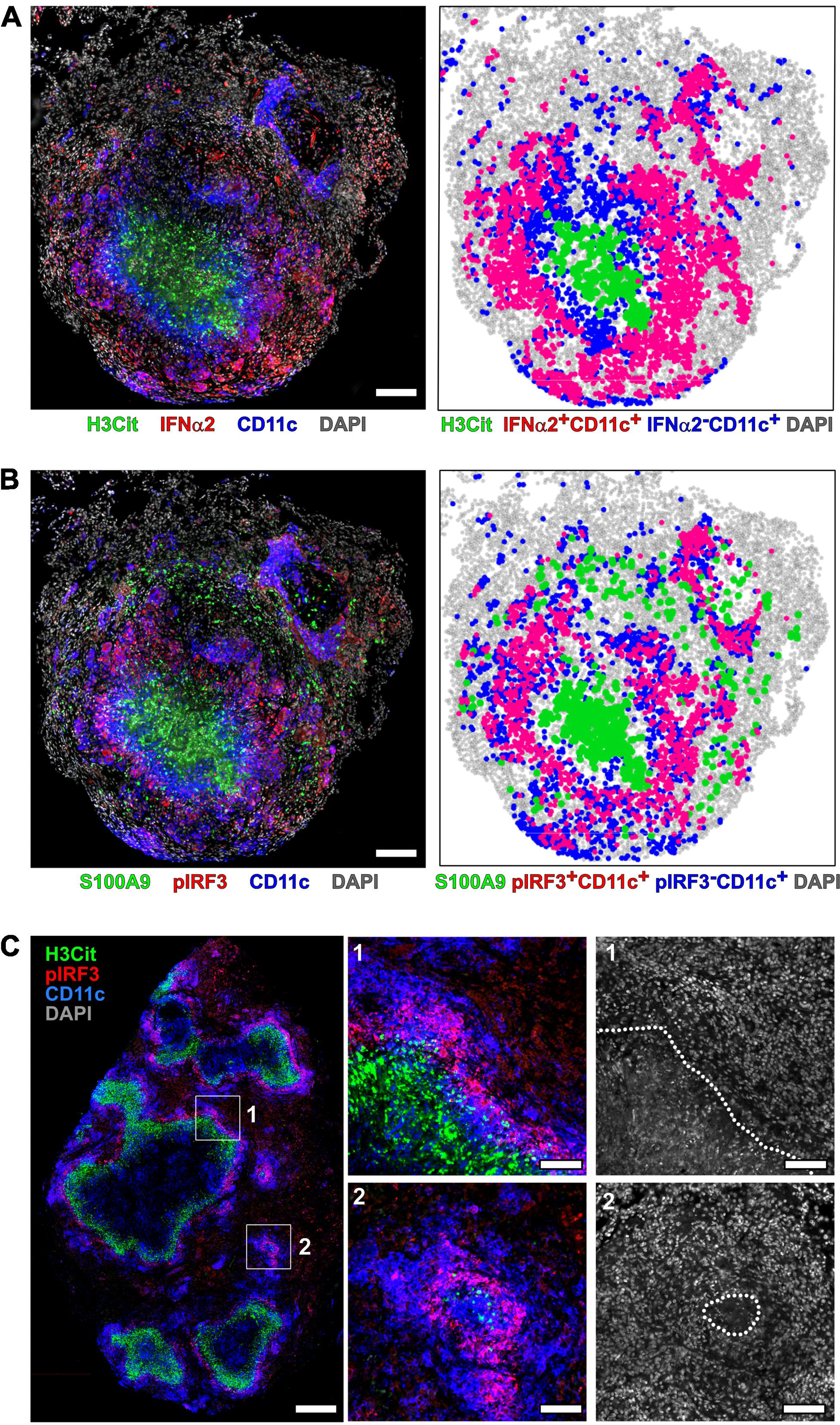
NETosis occurs in proximity to caseum and IFN**α**2-expressing epithelioid macrophages. **(A,B)** Cynomolgus macaque granulomas were stained for (A) IFNα2 (red), H3Cit (green), CD11c (blue, a marker for macrophages and dendritic cells), and nuclei (DAPI, gray) or (B) pIRF3 (red), S100A9 (green, neutrophil marker), CD11c (blue), and nuclei (DAPI, gray). Cells in the imaged sections (left) were segmented into subsets and plotted against the position of all the cells in the granuloma (grey) to highlight the spatial position of macrophages that may be expressing IFNα2 (red) in context with CD11c^+^ macrophages, H3Cit^+^ cells, and neutrophils (right panels). 20x magnification; scale bars represent 100 μm. H3Cit+ neutrophils (green) accumulate at the caseum-epithelioid macrophage interface in severely inflamed tissues (region 1) but are also present in small focal regions that may represent early sites of caseation (region 2). Dotted lines represent the border of regions containing cells with intact nuclei and degraded DNA that is indicative of caseation. 20x magnification; scale bar represents 500 μm in the whole tissue image (left) and 100 μm in the zoomed in regions (middle and right).

To better understand how NETosis and type 1 IFN expression are linked in lesions with poor immune control, we examined samples of TB pneumonia, a severe form of disease characterized by overwhelming inflammation and multiple caseous foci. Studying this lesion type gave us the ability to investigate how NETosis varied with IFNα2 expression in highly inflamed non-necrotic and necrotic regions (Figure 7C). As with necrotic granulomas, we found that NETotic neutrophils were only present at interface of pIRF3^+^ epithelioid macrophages and caseum (Figure 7C, region 1). We also observed small aggregates of pIRF3^+^ macrophages in non-necrotic regions and these regions contained H3Cit^+^ neutrophils (Figure 7C, region 2). When we examined the integrity of the nuclei in the center of these regions by comparing the DAPI staining across the field of view, we noted a loss of nuclear integrity in the center of these aggregates, indicating that these foci represented sites of developing necrosis. These observations suggest that neutrophils migrate to necrotic foci, even at the earliest stages of caseation, and these cells are exposed to type 1 IFN before they undergo NETosis. This suggests that the combination of macrophage type 1 IFN expression, neutrophil recruitment, and neutrophil NETosis contribute to development of caseous necrosis in primate granulomas. In addition, NETs can induce type I IFN expression^75, 76^, potentially providing evidence for a positive feedback loop involving neutrophils, type I IFN expression, and granuloma macrophages.

## DISCUSSION

There is a growing appreciation for the association of increased neutrophil abundance with active TB disease. However, it is still unknown if the presence of neutrophils in the lungs of active TB patients is consequential, or if the neutrophils are bystanders reacting to an uncontrolled infection. In particular, the details on how specific neutrophil responses and effector functions impact TB disease have remained elusive. We have used genetic and chemical approaches to show that a specific response by neutrophils, NETosis, directly contributes to *Mtb* replication and pathogenesis. Contrary to the antimicrobial nature associated with NETs, we find that *Mtb* survives exposure to NETs and can exploit NETosis to promote replication *in vitro*. This is particularly surprising given that neutrophil granule proteins that are released on NETs have been shown to inhibit *Mtb* replication when added to cultures of *Mtb* or *Mtb*-infected macrophages *in vitro*^77–80^. Therefore, the mechanisms by which Mtb defends itself against the antimicrobial effects of NETs and how NETosis promotes *Mtb* replication remain unknown. Many bacterial species secrete nucleases to utilize extracellular DNA as a nutrient source or to degrade NETs as a defense mechanism^81, 82^. *Mtb* has been shown to secrete a nuclease, Rv0888, that can degrade DNA and RNA^83^, but its role in *Mtb* pathogenesis has yet to be elucidated. The process of NETosis could also release other nutrients from neutrophils that promote *Mtb* growth. In addition, NETs and the proteins that accompany neutrophil degranulation are associated with lung damage^9, 84–86^ and in granulomas the collateral damage these factors cause to nearby cells, including macrophages, could inhibit antimicrobial functions and facilitate release of nutrients from bystander cells. Our studies highlighted a close association between NETosis and caseum in granulomas, specifically in a region that has previously been identified as harboring many bacteria^73^. Therefore, the contribution of NETosis to granuloma necrosis and caseation could be another mechanism by which NETosis promotes an environment for *Mtb* to thrive. Moreover, by entangling but not killing the *Mtb*, neutrophils undergoing NETosis could promote *Mtb* aggregation that protects the bacilli from environmental and antibiotic stress. Indeed, extracellular *Mtb* aggregates have been observed within the acellular rim of necrotic lesions^87–89^.

The process of caseation in granulomas is not well understood but our results showing that NETotic neutrophils are prominent in the smallest caseous foci we could detect suggests that neutrophilic infiltration and NETosis are consequential to the early formation of caseum. We were not able to determine if NETosis is followed by necrosis in adjacent macrophages, or if neutrophils are recruited to necrotic macrophages and then undergo NETosis, or if both scenarios occur simultaneously, but all these options may contribute to granuloma-level caseation. NETosis has been associated with macrophage death by pyroptosis in murine models of sepsis^90^, and further work investigating interactions between neutrophils and epithelioid macrophages in macaque granulomas may identify causal or temporal relationships between these behaviors and caseation. Considering the link between neutrophilic inflammation and TB pathology, the outcomes of neutrophil-driven caseation has implications for control of bacterial dissemination, especially in settings where necrotic granulomas have the potential to invade airways or blood vessels. NETosis may also contribute to the progression of necrotic granulomas into cavitary lesions, a process that promotes active TB disease and bacterial transmission. Taken together, these hypotheses support a paradigm that links excessive neutrophilic inflammation and NETosis-driven caseation with reduced control of TB at the lesion and systemic levels.

Our data highlights the association of NETosis with type I IFN production by epithelioid macrophages but does not rule out type I IFN from other cell types. Alveolar macrophages and plasmacytoid DCs (pDCs) also express type I IFN^91, 92^, and neutrophils are likely to encounter these cells as they are recruited into granulomas. In addition, our *in vitro* studies suggest that neutrophils themselves also produce sufficient type I IFN to induce NET release. Despite the well-established connection between type I IFN and NETosis, how type I IFN signaling contributes to NETosis remained elusive. We have discovered that type I IFN signaling in neutrophils functions downstream of histone citrullination to promote the formation of chromatin containing vesicles and their release as NETs. Our findings reveal that there are multiple points that regulate NETosis during *Mtb* infection while highlighting that histone citrullination can occur without efficient release of NETs. The specific role for type I IFN signaling during these later steps of NETosis could also be relevant to the other contexts where type I IFN promotes NETosis and exacerbates disease, such as systemic lupus erythematosus (SLE). In addition to this newly discovered neutrophil-intrinsic role for type I IFN signaling during NETosis, type I IFN signaling in non-neutrophils promotes histone citrullination in neutrophils in susceptible mice during *Mtb* infection^9^, highlighting another way that type I IFN can regulate NETosis. In this regard, the histone citrullination observed in neutrophils adjacent to IFNα2-expressing epithelioid macrophages likely results from IFN-regulated factors, and not necessarily type I IFN itself. In addition to a role for type I IFN signaling in promoting NETosis, NETs themselves can induce type I IFN production by pDCs and myeloid cells through toll like receptor 9 (TLR9) and STING-dependent signaling pathways^93–97^. Taken together, these features may contribute to a cycle of NET-driven immunopathology where NETs promote epithelioid macrophage type I IFN expression, and this activates a cascade of pathogenic neutrophil- and type I IFN-regulated responses in granulomas. A number of studies have linked increased and sustained levels of type I IFN signaling with TB pathology in mice and humans^14, 24, 98, 99^. Based on our data that NETosis promotes *Mtb* replication and pathogenesis, NETosis could contribute to the ways that type I IFN signaling impedes control of *Mtb* infection. In addition to impacting type I IFN signaling, NETs can modulate other myeloid cell activities, including cytokine production and phagocytosis^100, 101^. NETosis also promotes the formation of low density neutrophils^102^, which are associated with poor TB outcomes^103, 104^. Therefore, the interactions between NETotic neutrophils and other cell types within granulomas, and how these interaction shape outcomes in TB, are complex and require further dissection.

The alarming rise of drug-resistant TB cases has made it clear that we are not equipped to successfully battle the TB epidemic. There is growing interest in host-directed therapies (HDT) that would be effective against both drug-sensitive and drug-resistant *Mtb*. Herein, we identify NETosis as a potential HDT target and highlight points of regulation of this process during *Mtb* infection. Our findings could also be applied to other diseases where excessive neutrophilic inflammation and NETosis is linked to pathology. This includes thrombosis where NETs provide the scaffold and stimulus for thrombus formation^105^. NETosis is also a prominent feature of lung pathology in influenza^106, 107^, SIV infection^108^, and COVID-19^109^. Thus, better understanding of how NETosis is regulated in different contexts could have broad implications for human health.

## ACKNOWLEDGEMENTS

This work was supported by NIH grants R01 AI132697 and U19 AI142784, a Burroughs Wellcome Fund Investigators in the Pathogenesis of Infectious Disease Award, and the Philip and Sima Needleman Center for Autophagy Therapeutics and Research to C.L.S., Potts Memorial Foundation postdoctoral fellowship to R.L.K, Stephen I. Morse Fellowship to S.K.N., and NIH grant T32 AI007172 to E.M.N. Work at the University of Pittsburgh was supported by NIH grants R01 AI164970 and R21 AI167710, and funding from the Wellcome Leap Delta Tissue Program to J.T.M.. We are grateful for the assistance of Dr. Sanja Sviben, Dr. Praveen Krishnamoorthy, and Dr. James Fitzpatrick at the Washington University Center for Cellular Imaging (WUCCI) with the scanning electron microscopy and confocal microscopy studies, which is supported by Washington University School of Medicine, the Children’s Discovery Institute of Washington University and St. Louis Children’s Hospital (CDI-CORE-2015-505 and CDI-CORE-2019-813) and the Foundation for Barnes-Jewish Hospital (3770).

## AUTHOR CONTRIBUTIONS

The experiments were designed by C.S.C., J.T.M., and C.L.S.. The experiments were executed by C.S.C., R.L.K., E.M.N, S.K.N., D.S.L., P.T., and J.T.M., with assistance from A.S. and S.M.C.. W.B. assisted with TEM studies. C.S.C., J.T.M., and C.L.S. analyzed the data. D.K. bred and maintained the mouse colonies. The manuscript was written by C.S.C., J.T.M., and C.L.S. and all authors provided edits and comments on drafts.

## METHODS

### *Mtb* strains and bacterial cultures

*Mtb* Erdman expressing GFP (GFP-*Mtb*^12, 13^) was used in all *in vitro* experiments, wild-type *Mtb* HN878 strain (kindly provided by Dr. Selvakumar Subbian) was used for the mouse infections, and wild-type *Mtb* Erdman was used in the nonhuman primate studies. *Mtb* strains were cultured at 37°C in 7H9 (broth) or 7H11 (agar) (Difco) medium supplemented with 10% oleic acid/albumin/dextrose/catalase (OADC), 0.5% glycerol, and 0.05% Tween 80 (broth). Cultures of GFP-*Mtb* were grown in the presence of kanamycin (20 μg/ml) to ensure plasmid retention.

### Mice

Adult mice (age 7–15 weeks) of both sexes were used and mouse experiments were randomized. C57BL/6J (000664), *Padi4*^-/-^ (030315), and C3HeB/FeJ (000658) mice were all purchased from Jackson Laboratory and bred in pathogen-free barrier facilities at the Washington University in Saint Louis. *Ifnar1*^-/-^ mice were kindly provided by Drs. Ashley Steed, Wayne Yokoyama, and Bob Schreiber. No blinding was performed during animal experiments. All procedures involving animals were conducted following the National Institute of Health guidelines for housing and care of laboratory animals and performed in accordance with institutional regulations after protocol review and approval by the Institutional Animal Care and Use Committee of The Washington University in St. Louis School of Medicine. Washington University is registered as a research facility with the United States Department of Agriculture and is fully accredited by the American Association of Accreditation of Laboratory Animal Care. The Animal Welfare Assurance is on file with OPRR-NIH. All animals used in these experiments were subjected to no or minimal discomfort. All mice were euthanized by CO_2_ asphyxiation, which is approved by The Panel on Euthanasia of the American Veterinary Association.

### Neutrophil isolation from mice

Bone marrow neutrophils were obtained using a single-step Percoll gradient^110^. Briefly, murine bone marrow was flushed out of the tibia and the femur in HBSS supplemented with 20 mM HEPES (Sigma) using a 23G needle (McKesson) and passed through a 70 µm cell strainer (CellTreat) after hypertonic lysis with of erythrocytes using a 0.2% NaCl solution. Cells were pelleted and the entire bone marrow was resuspended in HBSS without CaCl_2_ and MgCl_2_, 20 mM HEPES (Sigma), and 0.5% HI FBS (Gibco).

Neutrophils were purified by centrifugation for 30 min at 1,300xg on a discontinuous Percoll gradient (GE Healthcare #17-0891-01) consisting of 62% (v/v) Percoll in HBSS, 20 mM HEPES (Sigma), and 0.5% HI FBS (Gibco). Neutrophils were recovered from the bottom of the tube.

### *In vitro* infection of neutrophils

Isolated bone marrow neutrophils were resuspended in HBSS and counted. 0.5x10^6^ cells were transferred to a 24 well plate containing glass coverslips and RPMI 1640 supplemented with 10% Heat inactivated FBS and incubated for 45 minutes. Logarithmically growing GFP-*Mtb* was counted after washing and sonicating in PBS to separate clumps of bacteria. *Mtb* were opsonized for 1h with 5% normal mouse serum (NMS) in RPMI before adding to the neutrophil cultures in the 24 well plate at an MOI of 20. Plates were centrifuged at 200xg for 10 minutes. For microscopy experiments, cells plus bacteria were incubated for 30 minutes at 37°C and 5% CO_2_ following centrifugation before removing all non-phagocytoses bacteria and replacing the media with RPMI 1640 supplemented with 10% Heat inactivated FBS, 1 mM CaCl_2_ and 1mM MgCl_2_. For infections used to monitor bacterial burden, non-phagocytosed bacteria were not removed.

### Fluorescent microscopy and quantification

For fluorescent microscopy of neutrophils *in vitro*, neutrophils on coverslips were fixed with 4% paraformaldehyde (PFA) and staining was performed with antibodies specific for H3Cit (citrulline R2 + R8 + R17, Abcam Ab5103, 1:200), myeloperoxidase (MPO, goat polyclonal R&D Systems, 1:200) or Ly6G (Clone 1A8, 1:200). DNA was labeled with 5 μg/ml Hoechst 33342 (Molecular Probes). To detect cells with compromised plasma membrane integrity indicative of cell death, neutrophils on cover slips were treated with Zombie NIR (BioLegend, 1:1000) for 5 min before PFA fixation and staining with Hoechst and other antibodies. A Nikon A1Rsi confocal microscope coupled with NIS software was used to take Z-stack images with a 60x oil immersion objective. ND2 files of Z-Stack confocal images were merged using FIJI software. Fields that were imaged and used for quantification were selected randomly based on areas containing around 100 cells to keep the cell number consistent between groups or treatments. Ridge detection analysis was performed on 8-bit images. Structures above the threshold described in supplementary figure S1 were counted and measured.

### Scanning electron microscopy (SEM)

Cultures on glass coverslips were washed with 0.15M cacodylate buffer warmed to 37°C and fixed overnight in a solution containing 2.5% glutaraldehyde and 2% PFA in 0.15 M cacodylate buffer at pH 7.4 that had been warmed to 37°C. Coverslips were then rinsed in 0.15 M cacodylate buffer 3 times for 10 minutes each and subjected to a secondary fixation in 1% osmium tetroxide in cacodylate buffer for one hour. Samples were then rinsed in ultrapure water 3 times for 10 minutes each and dehydrated in a graded ethanol series (30%, 50%, 70%, 90%, 100% x3) for 10 minutes each. Once dehydrated, samples were loaded into a critical point drier (Leica EM CPD 300, Vienna, Austria) that was set to perform 12 CO_2_ exchanges at the slowest speed. Samples were then mounted on aluminum stubs with carbon adhesive tabs and coated with 5 nm of carbon and 6 nm of iridium (Leica ACE 600, Vienna, Austria). SEM images were acquired on an FE-SEM (Zeiss Merlin, Oberkochen, Germany).

### Transmission electron microscopy (TEM)

TEM of neutrophils has been described previously^54^. Neutrophils either mock-infected or infected with *Mtb* were released from the coverslips using 10mM EDTA for 5 min and collected into a tube and pelleted. For ultrastructural analysis, samples were fixed in 2% PFA/2.5% glutaraldehyde in 100 mM sodium cacodylate buffer for 2 h at room temperature and then overnight at 4°C. Samples were washed in sodium cacodylate buffer, embedded in 2.5% agarose, and postfixed in 2% osmium tetroxide (Ted Pella Inc., Redding, CA) for 1 h at room temperature. After three washes in dH_2_O, samples were en bloc stained in 1% aqueous uranyl acetate (Electron Microscopy Sciences, Hatfield, PA) for 1 h. Samples were then rinsed in dH_2_0, dehydrated in a graded series of ethanol, and embedded in Eponate 12 resin (Ted Pella Inc). Sections of 95 nm were cut with a Leica Ultracut UCT ultramicrotome (Leica Microsystems Inc., Bannockburn, IL), stained with uranyl acetate and lead citrate, and viewed on a JEOL 1200 EX transmission electron microscope (JEOL USA Inc., Peabody, MA) equipped with an AMT 8-megapixel digital camera and AMT Image Capture Engine V602 software (Advanced Microscopy Techniques, Woburn, MA).

### Plating for CFU from *Mtb*-infected neutrophil cultures

After 48 hours of incubation, neutrophils were lysed by addition of a final concentration of 0.5% Triton X-100 (Sigma) directly into the medium. Lysates were 10-fold serially diluted in 0.05% Tween 80 in PBS and plated on Middlebrook 7H11 agar plates. After 3-4 weeks at 37°C in 5% CO_2_, colonies were counted and total cell numbers were calculated.

### *Mtb* infection of mice

*Mtb* infection of mice have been previously described^11^. Briefly, before infection, exponentially replicating *Mtb* HN878 strain were washed in PBS + 0.05% Tween 80 and sonicated to disperse clumps. 7- to 15-week-old female mice were exposed to 8 × 10^7^ CFU of *Mtb* in an Inhalation Exposure System (Glas-Col), which delivers ∼100 bacteria to the lung per animal. At 24 hours post infection, the bacterial titers in the lungs of at least two mice were determined to confirm the dose of *Mtb* inoculation. The dose determined from these mice is assumed to represent the average dose received by all mice in the same infection. Bacterial burden was determined by plating serial dilutions of organ homogenates onto 7H11 agar plates. Plates were incubated at 37°C in 5% CO2 for 3 weeks prior to counting colonies.

### Histology of mouse lung tissue

Lung tissue fixed in 4% PFA were dehydrated using sucrose gradient and mounted into blocks using Fisher OCT in liquid nitrogen. Frozen lung sections were cut at 5 μm. Antigen retrieval was performed for 5 min using 1% SDS. Samples were permeabilized with 0.5% Triton X-100 in PBS containing 1% BSA for 5 min, blocked for 1 h using 5% donkey serum and then incubated with a rabbit monoclonal H3Cit antibody (citrulline R2 + R8 + R17, clone EPR20358-120; Abcam) at 1:500 dilution overnight in a humidified chamber at 4°C. Samples were washed with PBS and incubated with a Alexa Fluor 555 conjugated secondary antibody (Invitrogen, 1:200), Alexa Fluor 488 conjugated MPO (Abcam, 1:200), Alexa Fluor 647 conjugated Ly6G antibody (BioLegend, 1:200), and Hoechst (1:1000) for 1 h at room temperature. Samples were mounted after removing tissue autofluorescence using Vector® TrueVIEW® Autofluorescence Quenching Kit. Z-stack images of large 8x8 area under 10x objective were taken using a Nikon A1Rsi confocal microscope coupled with NIS software, and mosaics were made with 20% overlap to cover the entire lung section.

### Flow Cytometry

Lungs were perfused with sterile PBS and digested at 37°C for 1 h with 625 μg/mL collagenase D (Roche) and 75 U/mL DNase I (Sigma). Single cell suspensions stained in PBS + 2% FBS in the presence of Fc receptor blocking antibody (BD Pharmingen) and stained with the antibodies against the following mouse proteins: SiglecF (clone E50-2440; BD Pharmingen), Ly6G (clone 1A8; BioLegend), CD11c (clone N418; BioLegend), CD11b (clone M1/70; BioLegend), MHCII (clone M5/114.15.2; BioLegend), CD45 (clone 30-F11; BioLegend), Ly6C (clone HK1.4; BioLegend), CD4 (clone RM4-5; BioLegend), TCRβ (clone H57-567; BioLegend), CD8a (clone 53-6.7; BioLegend), CD19 (clone 6D5; BioLegend). Cells were stained for 20 minutes at 4°C and then fixed in 4% PFA (Electron Microscopy Sciences) for 20 minutes at room temperature. For intracellular H3Cit staining, PFA fixed cells were permeabilized with BD perm/wash buffer for 5 min. Cells were blocked with 2% BSA in PBS and then incubated with anti- H3Cit antibody (citrulline R2 + R8 + R17, clone EPR20358-120; Abcam) overnight at 4°C. After incubation, cells were washed with PBS and incubated with Alexa Fluor 647 conjugated secondary antibody (Invitrogen,1:200) and anti-Ly6G antibody (clone 1A8; BioLegend) for 1h at RT. Following incubation, cells were washed with PBS and resuspended in FACS buffer. Flow cytometry was performed on a Cytek Aurora (Cytek Biosciences). Spectral unmixing was performed using Cytek Aurora software and data was analyzed with FlowJo (Tree Star Inc.). Gating strategies in Figure S7.

### Immunofluorescence on nonhuman primate granulomas

Immunofluorescence was performed on granulomas from *Mtb*-infected cynomolgus macaques (*Macaca fascicularis*) involved in completed studies at the University of Pittsburgh that were approved by the University of Pittsburgh’s IACUC. Animals used in this study, by animal identifier and duration of Mtb infection (days post infection; dpi), include 9409 (69 dpi), 907 (76 dpi), 18214 (84 dpi), 18314 (91 dpi), 1707 (94 dpi), 18514 (118 dpi), 18414 (125 dpi), 6409 (140 dpi), 4710 (254 dpi), 12603 (277 dpi), 9905 (464 dpi). All animals were infected with Erdman-strain *Mtb* via bronchoscopic instillation except for monkey 4710, which was infected via aerosol route, and all were monitored as previously indicated^111, 112^ for the study’s duration. At the end of the study, the animals were all assessed to have active TB based on clinical and radiographic features. Details for each animal are indicated in Table S1. Lung granulomas were harvested from each animal at the time of necropsy and fixed in 10% neutral buffered formalin and embedded in paraffin. 5-μm thick sections were cut from these blocks and used for staining. The samples included here were selected based on the presence of granulomas and some sections contained multiple granulomas, thus bacterial burdens per granuloma were not available for most of these lesions. Immunofluorescent staining was performed as previously indicated^19, 21, 72^ for antigens including anti- S100A9/calprotectin (clone MAC387; Thermo Fisher Scientific, Waltham, MA), anti-histone H3 (citrullinated at R2+R8+R11, clone 11D3, Caymen Chemical, Ann Arbor, MI), anti-CD11c (clone 5D11; Leica Biosystems, Deer Park, IL), and anti-IFNα2 (rabbit polyclonal; Thermo Fisher Scientific). Species- and isotype-specific secondary antibodies were purchased from Jackson ImmunoResearch (West Grove, PA) or Thermo Fisher Scientific. Where anti-S100A9 and anti-H3Cit were used in the same section, H3Cit was stained with IgG1-specific secondary antibodies and S100A9 was labeled with Thermo Fisher Scientific’s Xenon labeling kit and used as a tertiary stain. A second tissue section was stained in parallel and used as an isotype control with the same secondary cocktail as a control to ensure the specificity of the staining. Coverslips were mounted on slides using ProLong Gold Mounting Media with DAPI (Thermo Fisher). 14-bit images of stained granulomas were acquired at 20x magnification on an e1000 epifluorescence microscope (Nikon, Melville, NY) using a DS-Qi2 camera (Nikon) and individual frames were stitched together at acquisition by Nikon Elements AR v4.50.

Image analysis was done on the ND2 image files with segmentation performed by QuPath version 0.3.2^113^ and graphical representations of cell phenotypes were prepared in CytoMAP version 1.4.21^114^.

### Statistical analysis for biological experiments

All data are from at least two independent experiments. Samples represent biological (not technical) replicates. No blinding was performed during animal experiments.

Statistical analyses were performed with Prism (v9.4.1; GraphPad Software) using unpaired two-tailed Student’s t tests, after the data were tested for normality, to compare between two conditions or one-way ANOVA with Tukey’s correction to perform multiple comparisons. Sample sizes were sufficient to detect differences as small as 10% using the statistical methods described. When used, center values and error bars represent means ± SEM. *P* < 0.05 was considered significant. *P* >0.05 was denoted *, ** for *P* < 0.01, *** *P* < 0.001, and **** *P* < 0.0001. Only significant differences are noted in the figures.

