## Supplemental Table 1 for "Type I IFN signaling mediates NET release to promote *Mycobacterium tuberculosis* replication and granuloma caseation"

| **Monkey** | **Sex** | **Age** | **Infection dose** | **Infection route** | **Infection duration (days)** | **PMID^a^** |
| --- | --- | --- | --- | --- | --- | --- |
| 907 | male | 9y 2mo | no data | instilled | 76 | 23749634 |
| 18214 | male | 8y 3mo | 39 | instilled | 84 | 28592427 |
| 18314 | male | 8y 3mo | 39 | instilled | 91 | 28592427 |
| 1707 | male | 9y 9mo | 10 | instilled | 94 | 23749634 |
| 18514 | male | 8y 3mo | 38 | instilled | 118 | 34255474 |
| 18414 | male | 8y 8mo | 38 | instilled | 125 | 34255474 |
| 6409 | male | 6y 8mo | no data | instilled | 140 | 24336248 |
| 4710 | male | 6y 9mo | 16 | aerosol | 254 | 35128020 |
| 12603 | male | 6y 4mo | no data | instilled | 277 | 19620341 |
| 9905 | female | 7y 11mo | no data | instilled | 464 | 19620341 |

Table S1. Demographic data for cynomolgus macaques from which lung samples were stained in histology experiments. ^a^Indicates a publication where data on an animal has been previously published.
