## Supplemental Figures for "Type I IFN signaling mediates NET release to promote *Mycobacterium tuberculosis* replication and granuloma caseation"

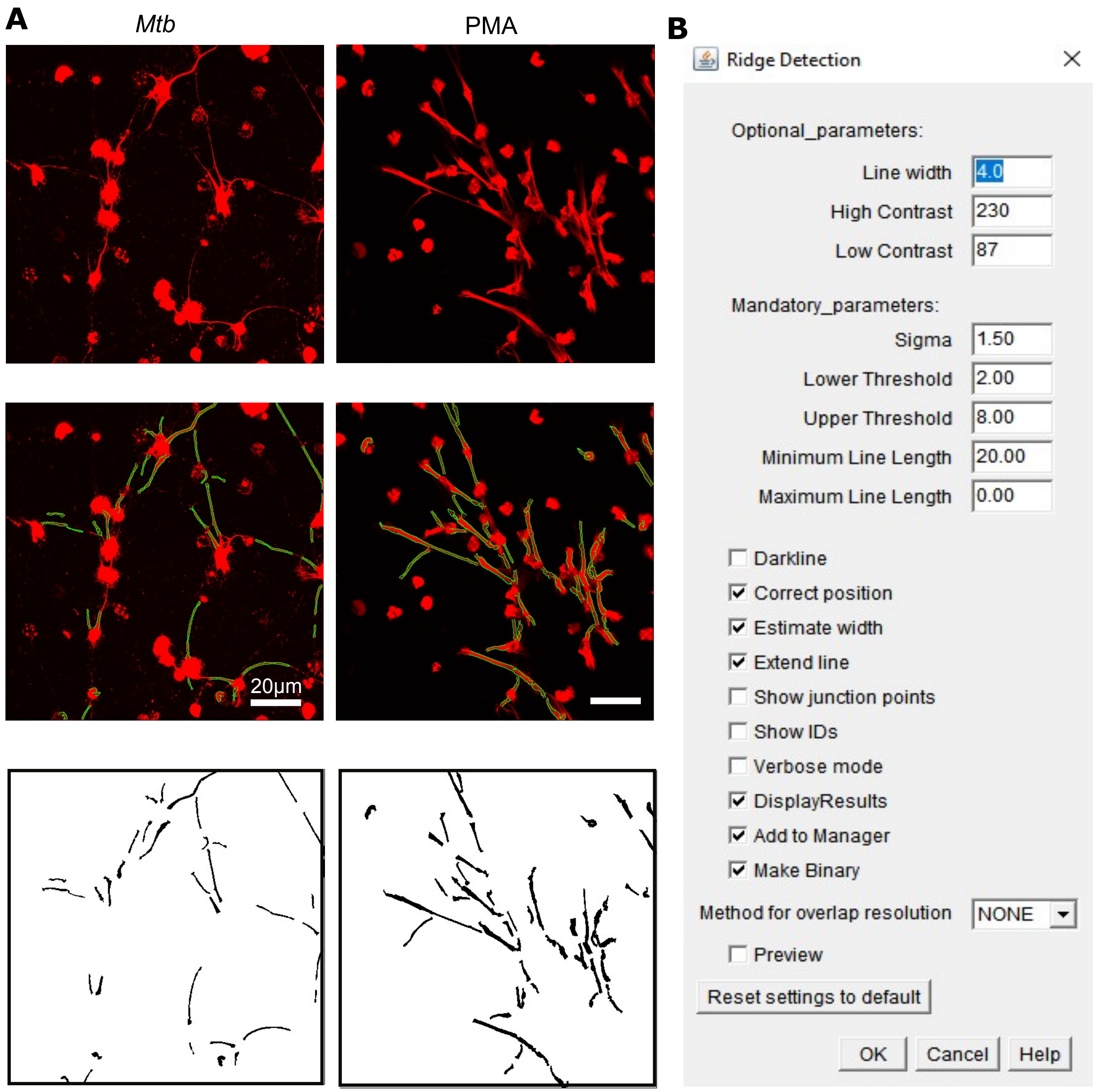


**Supplemental Figure 1. Method for quantification of extracellular web like structures (NETs).**
**(A)** Representative images showing analysis to quantify H3Cit$^{+}$ webs in confocal Z-stack images.
**(B)** Parameters followed using the Ridge Detection algorithm of FIJI software to quantify webs of NETs.

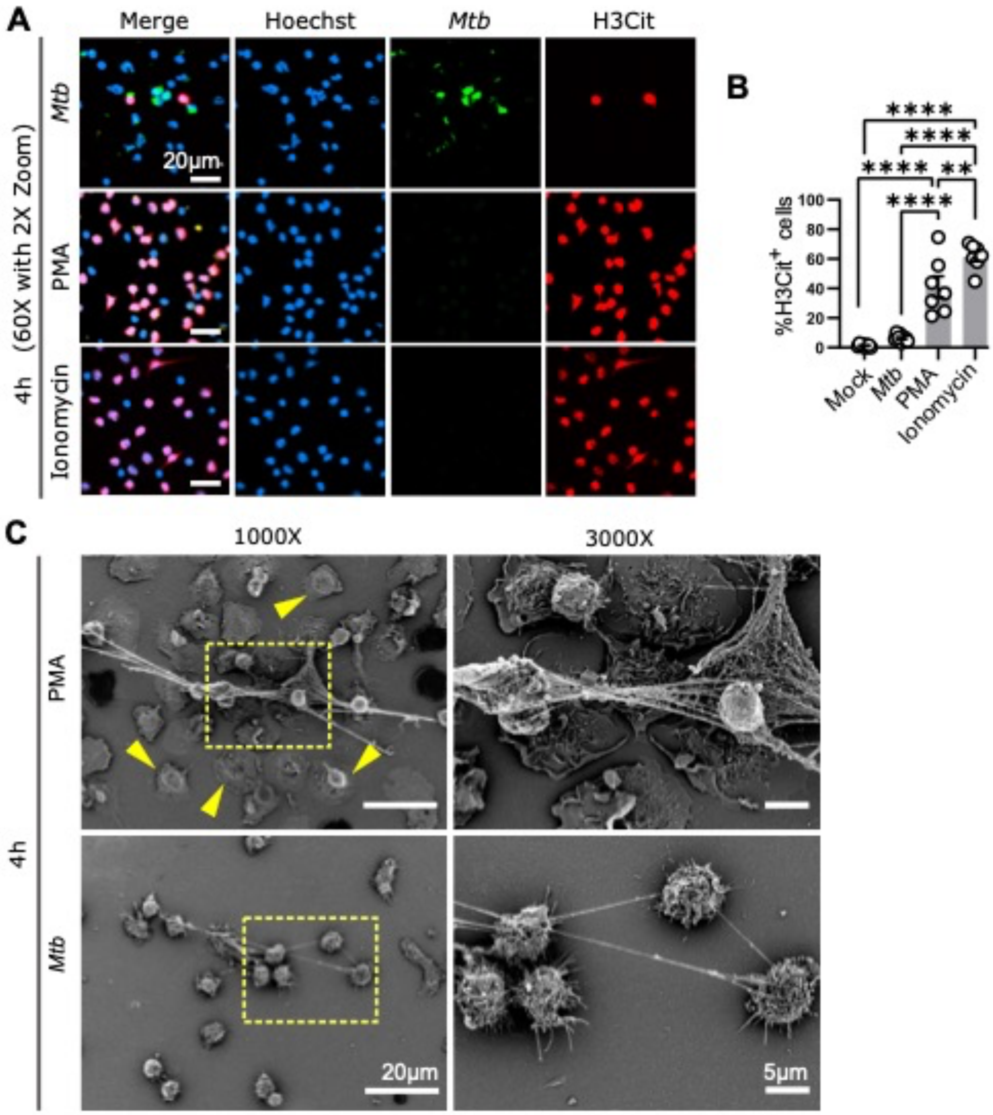


**Supplemental Figure 2. Neutrophil responses 4 hours following *Mtb* infection or treatment with PMA or ionomycin**

**(A)** Representative confocal images showing bone marrow neutrophils from C57BL/6J mice infected with GFP-*Mtb* at an MOI 20 or treated with PMA (100nM) or ionomycin (5μM) for 4h and stained for citrullinated histone 3 (H3Cit) and DNA (Hoechst). GFP-*Mtb* is also shown.

**(B)** The percentage of Hoechst$^{+}$ cells that were also H3Cit$^{+}$ per field under 60x objective were quantified using ImageJ software and plotted. Each datapoint represents data from one field and a minimum of 3 fields containing 20-200 cells/area for each condition were quantified. Bar graph of data represents mean ± SEM. $^{**}P < 0.01$; $^{****}P < 0.0001$ by one-way ANOVA with Tukey's correction.

**(C)** Representative scanning electron microscopy showing neutrophil morphology after 4 h coculture with *Mtb* or treatment with PMA. Yellow arrows denote the flattening of neutrophils in the presence of PMA. Clustered release of chromatin was observed during PMA treatment compared to thread like NETs released during *Mtb* infection. Images on the right are zoomed in from the region in the yellow box.

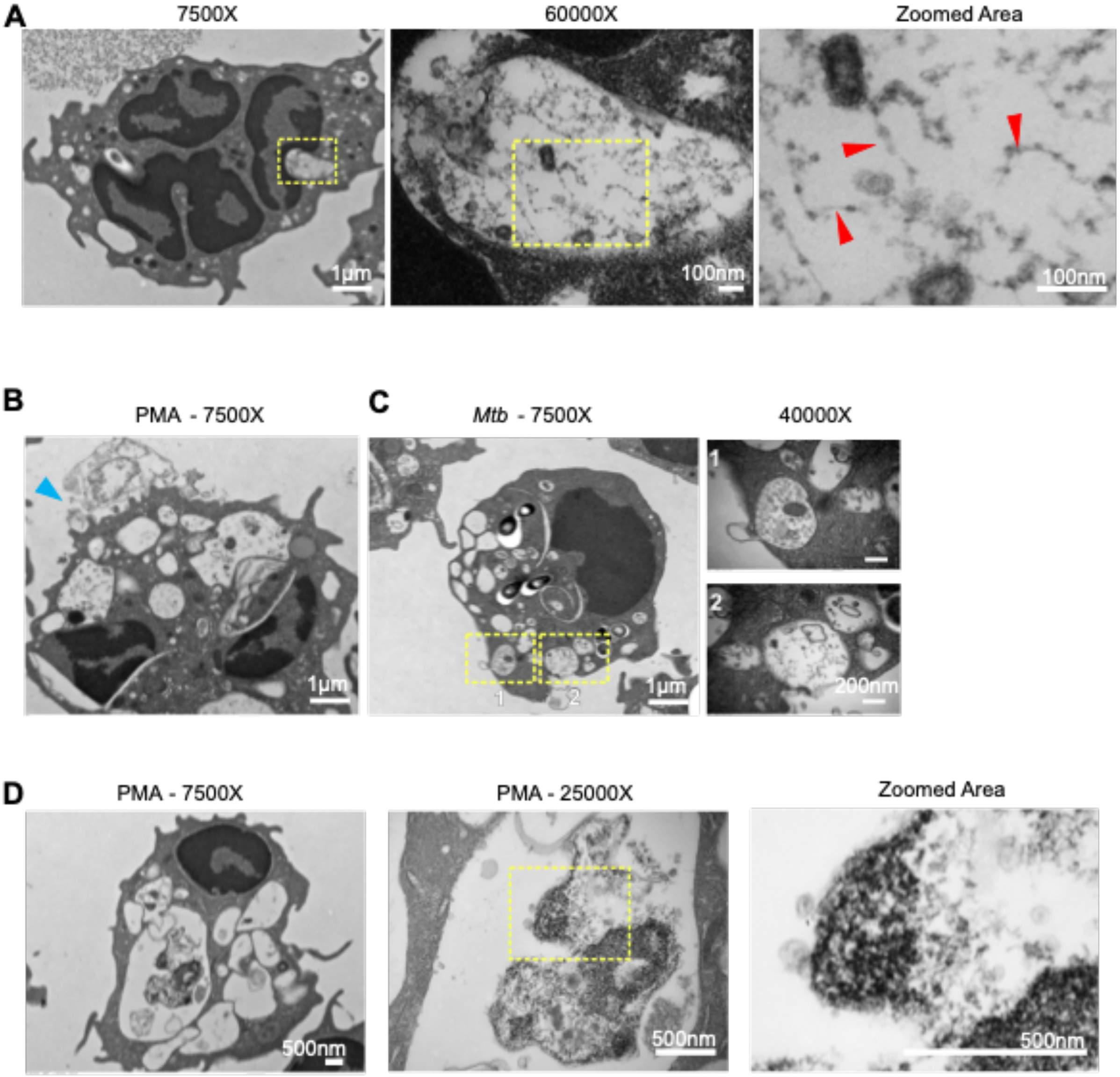


**Supplemental Figure 3. TEM of cellular processes observed during NETosis**

**(A)** Representative TEM images of WT neutrophils infected with *Mtb* at an MOI of 20 for 4h. The region in the yellow box is sequentially magnified in the right panels to highlight the “beads on a string” DNA and histone structures (red arrows) within vesicles (decondensed chromatin vesicles).

**(B)** Representative TEM images of neutrophils after 4 hours of treatment with PMA-showing release of vesicles, possibly through rupture of outer membrane (blue arrow).

**(C)** Representative TEM images of *Mtb*-induced release of vesicles, possibly through pores or membrane fusion, highlighted by the magnification of the areas in the yellow boxes.

**(D)** Representative TEM images of PMA-induced vesicles containing electron dense material seeming to unwind to decondensed “beads on a string” chromatin, highlighted by the magnification of the areas in the yellow boxes.

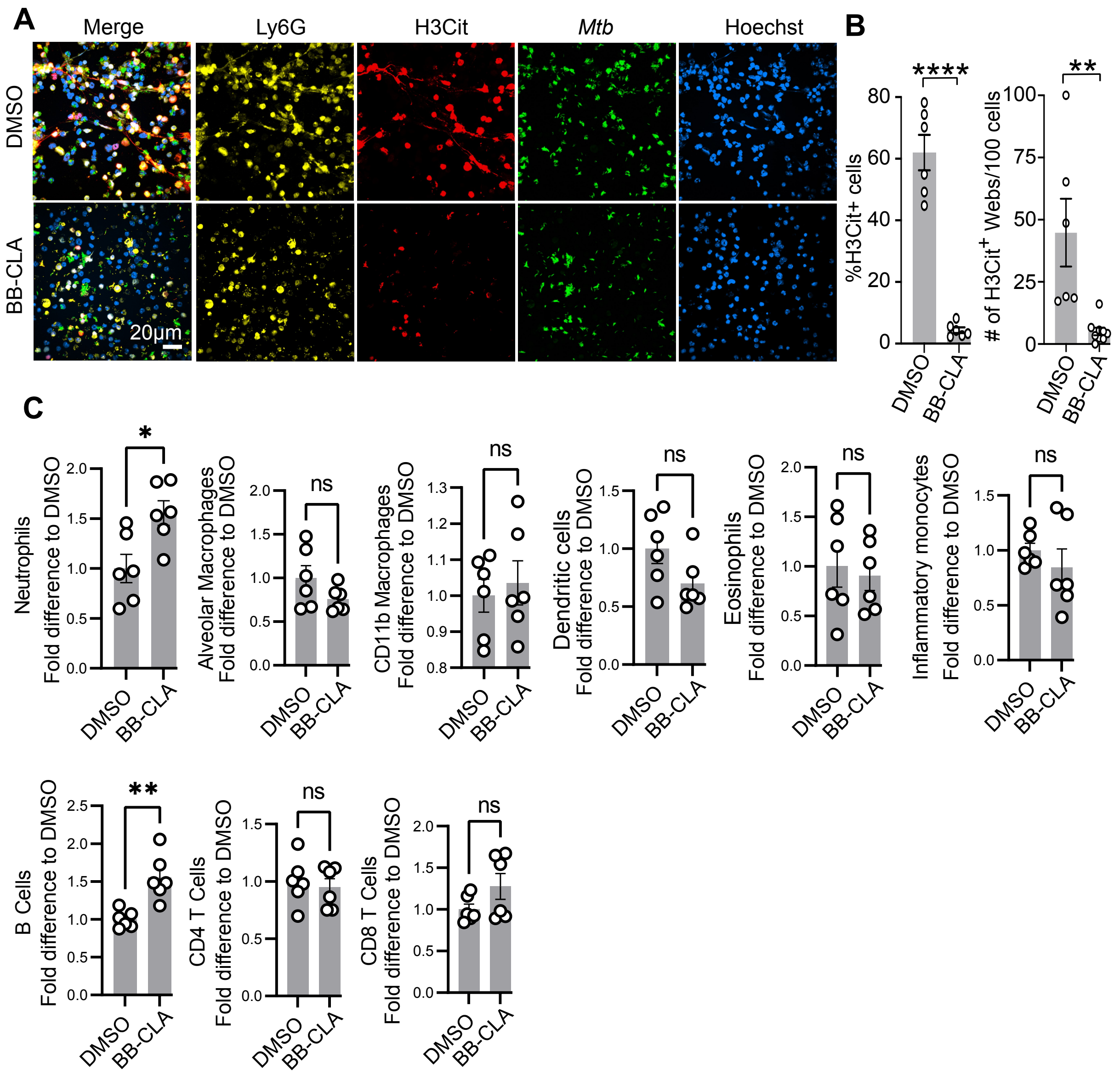


**Supplemental Figure 4. Effects of BB-Cl-amidine *in vitro* and *in vivo* during *Mtb* infection.**

**(A)** Representative confocal images showing immunofluorescence staining of neutrophils from WT C57BL/6J mice infected with GFP-*Mtb* at an MOI of 20 and treated with BB-Cl-amidine (BB-CLA, 50 µM) or DMSO for 18 h and stained for citrullinated histone 3 (H3Cit), a neutrophil marker (Ly6G), and DNA (Hoechst). GFP-*Mtb* is also shown.

**(B)** The percentage of Hoechst$^+$ cells that were also H3Cit$^+$ (left) and the number of extracellular H3Cit$^+$ webs per 100 Hoechst$^+$ nuclei (right) per field under 60x objective were quantified using ImageJ software and plotted. Each datapoint represents a single field and a minimum of 6 fields containing 20-200 cells/area were quantified from 2 independent experiments. Bar graph of data represents mean ± SEM. **P < 0.01 and ****P < 0.0001 by unpaired t test.

**(C)** Each datapoint is the frequency of CD45$^+$ cells for that cell type within a given mouse expressed relative to the average frequency in mice treated with DMSO vehicle, where at least 5 mice for each treatment were used in two independent experiments. Bar graph of data represents mean ± SEM. *P < 0.05; **P < 0.01; ns denotes not significant by or unpaired t-test.

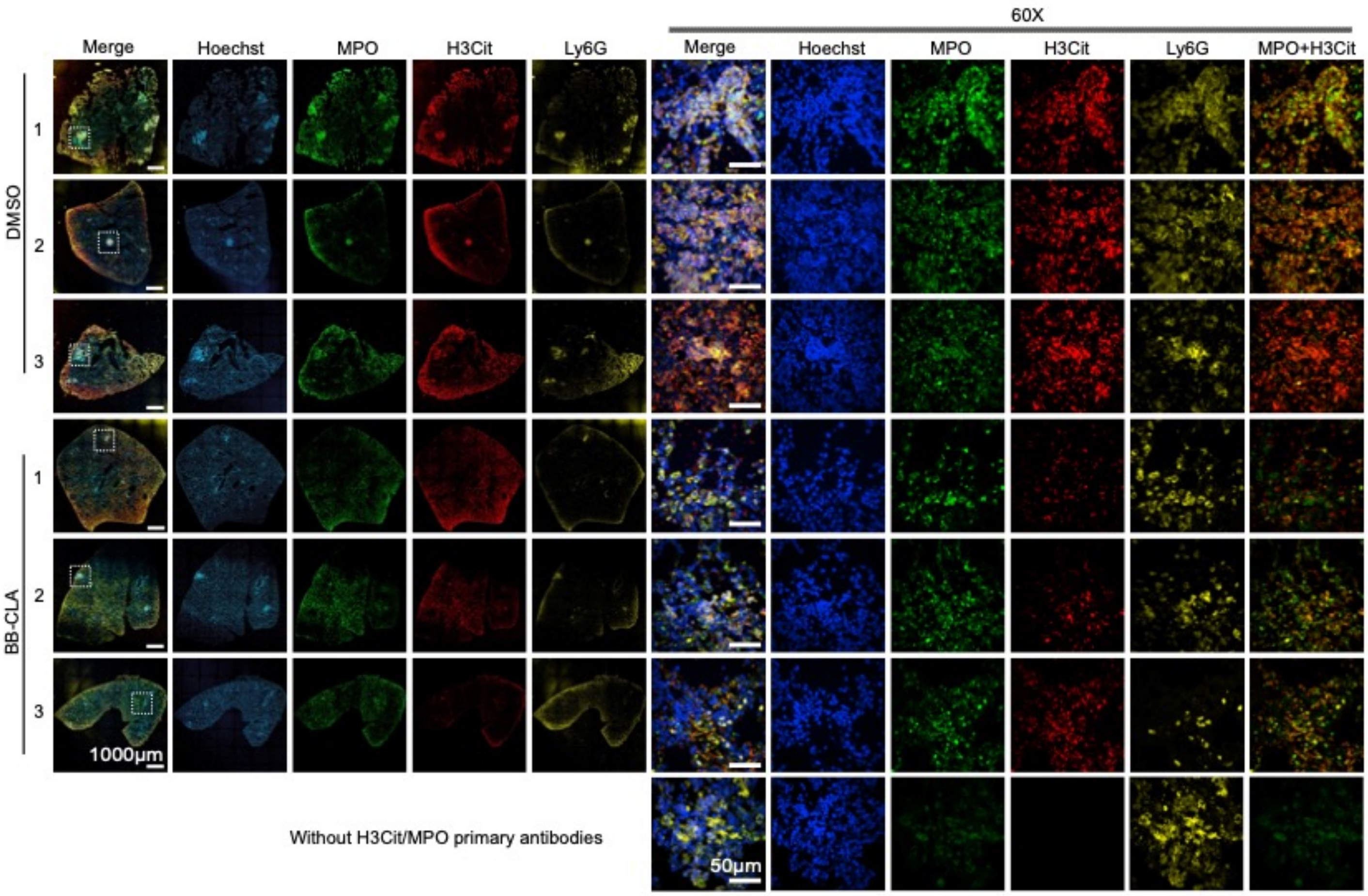


**Supplemental Figure 5. Representative lung histology samples from individual C3HeB/FeJ mice.** Representative images showing immunofluorescence staining of lung sections from *Mtb*-infected C3HeB/FeJ mice at 21 dpi that were probed with antibodies to detect citrullinated histone H3 (H3Cit; red), MPO (green, a marker for neutrophil granules), Ly6G (yellow, a marker for neutrophils) and DNA (Hoechst, blue). The entire lung section is shown along with a 60x zoomed in region denoted by the white box.

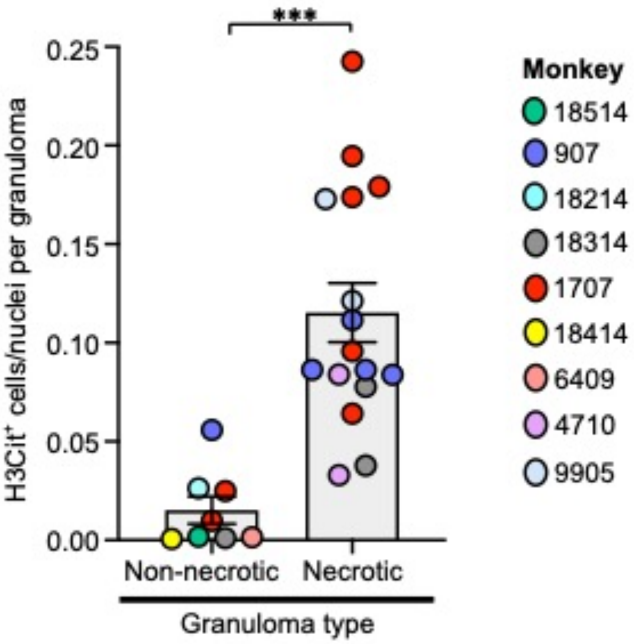


**Supplemental Figure 6. NETosis is observed more often in necrotic macaque granulomas than non-necrotic granulomas.**

Image analysis was performed on granulomas from macaques with active TB to quantify H3Cit$^+$ cells per granuloma cross section and the overall number of cells per granuloma was determined by quantifying the total number of nuclei per granuloma. Granulomas were classified as non-necrotic or necrotic according to the absence or presence of degraded nuclear DNA in the section, respectively, where granulomas containing diffusely-stained DNA that is indicative of caseation were considered to be necrotic. The normalized frequencies of H3Cit$^+$ cells in non-necrotic and necrotic granulomas were compared, showing that H3Cit$^+$ cells were significantly more abundant in necrotic granulomas compared to non-necrotic granulomas. Each point represents an individual granuloma and each color represents an animal, with the animals listed in ascending order according to their length of time post-infection before necropsy. Bar graph of data represents mean ± SEM and *** indicates $P<0.001$ by unpaired t test.

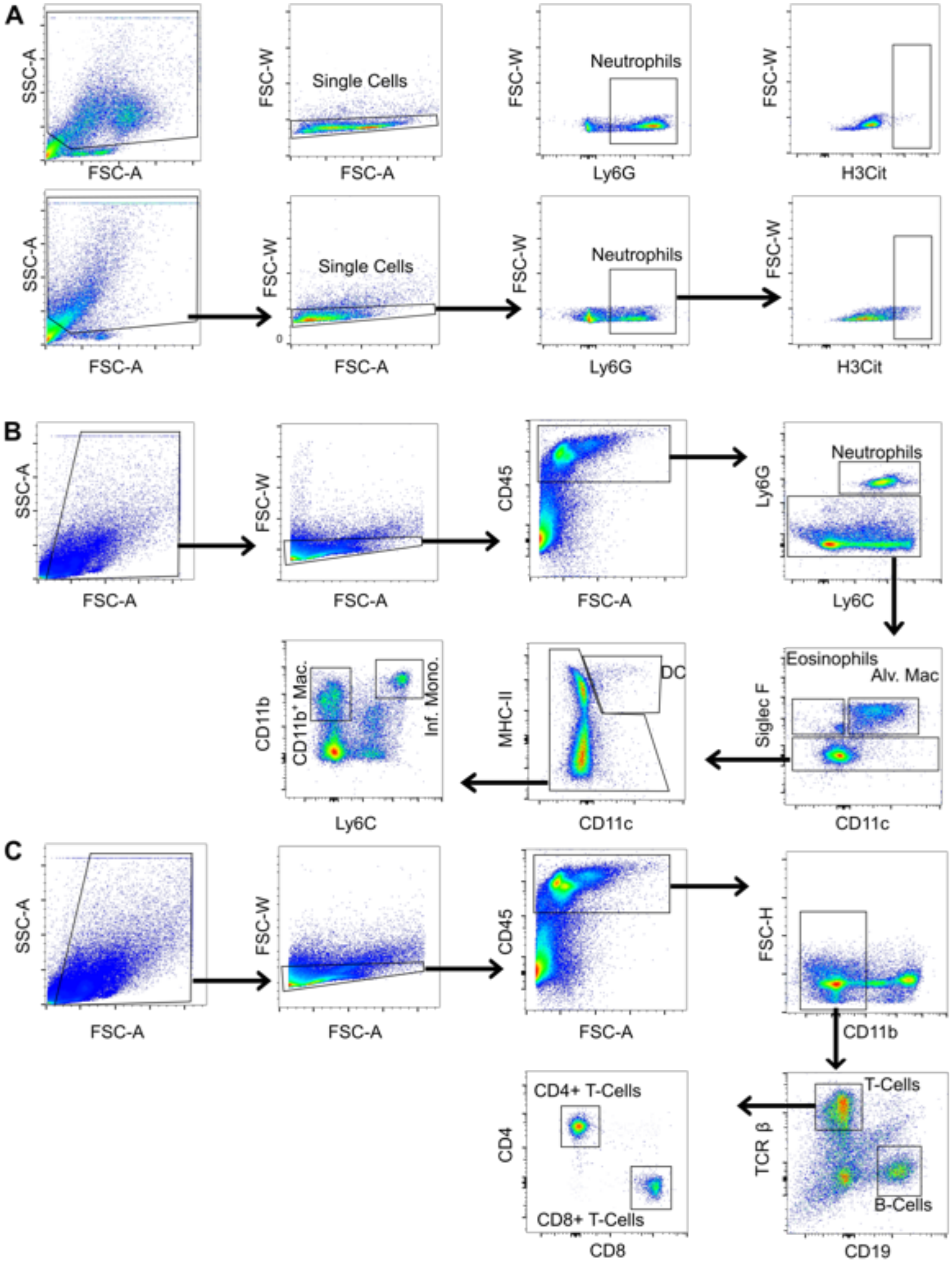


**Supplemental Figure 7. Gating Strategy for Flow Cytometry.**
**(A)** Identification of H3Cit$^+$ neutrophils from *in vitro* cultures at 4 hpi (top) and 18 hpi (bottom).
**(B)** Identification of innate immune cell populations from *Mtb*-infected murine lungs.
**(C)** Identification of T and B cell populations from *Mtb*-infected murine lungs.
